# Differential Activity of MAPK signalling Defines Fibroblast Subtypes in Pancreatic Cancer

**DOI:** 10.1101/2023.10.30.564752

**Authors:** Lisa Veghini, Davide Pasini, Pietro Delfino, Rui Fang, Dea Filippini, Christian Neander, Caterina Vicentini, Elena Fiorini, Francesca Lupo, Sabrina L. D’Agosto, Carmine Carbone, Michele Bevere, Diana Behrens, Claudio Luchini, Rita T. Lawlor, Aldo Scarpa, Giulia Biffi, Phyllis F. Cheung, Jens T. Siveke, Vincenzo Corbo

## Abstract

Fibroblast heterogeneity is increasingly recognised across cancer conditions. Given their important contribution to disease progression, mapping out fibroblasts’ heterogeneity is critical to devise effective anti-cancer therapies. Cancer-associated fibroblasts (CAFs) represent the most abundant cell population in pancreatic ductal adenocarcinoma (PDAC). Whether CAF phenotypes are differently specified by PDAC cell lineages remains to be elucidated. Here, we reveal an important role for the MAPK signalling pathway in the definition of PDAC CAF phenotypes. We identify the myCAF transcriptional phenotype as uniquely dependent on proficient MAPK signalling. In addition, CAFs displaying elevated MAPK activity are specifically anchored to basal-like/squamous PDAC cells and define tumour subdomains with reduced frequency of CD8+ T cells. We characterize the single-cell transcriptome of mouse PDAC tumours in response to MAPK inhibition and identify gene expression signatures of MAPK^high^ CAFs, which suggest immunoregulatory functions. Accordingly, a gene expression signature of MAPK^high^ CAFs correlates with poor prognosis in several human cancer conditions, including PDAC, and with reduced response to immune checkpoint inhibition in immune-reactive solid tumours. Altogether, our data expand our knowledge on CAF phenotype heterogeneity and reveal a new strategy for targeting of myofibroblastic CAFs *in vivo*.

## INTRODUCTION

Fibroblasts functionally contribute to disease progression and therapy response in solid tumours ^1,2^. The diversity of phenotypes and functions displayed by cancer-associated fibroblasts (CAFs) within the tumour microenvironment (TME) is dependent on their cellular origin, spatial localization, and disease context ^2,3^. In pancreatic ductal adenocarcinoma (PDAC), different subtypes and specialized functions of CAFs have been reported ^4–10^. Myofibroblastic CAFs (myCAFs) and inflammatory CAFs (iCAFs) are two distinct states consistently reported in the PDAC TME ^4,6,8^ for which mechanisms of cancer-induced rewiring have been provided ^4^. The failure of agnostic targeting of PDAC CAFs in preclinical and clinical settings ^11–14^ demonstrates our current insufficient understanding of fibroblast heterogeneity to properly guide therapeutic strategies. Here, we hypothesized that different neoplastic cell states contribute to PDAC fibroblast heterogeneity. Different transcriptional states of PDAC malignant cells have been described^15–19^, which result from the integration of cell intrinsic and extrinsic inputs ^20–22^. Transcriptional infidelity to the pancreatic endoderm and acquisition of exogenous gene programs (e.g., squamous/basal-like) is invariably associated with a more aggressive biological behaviour of cancer cells ^23^. However, we have a limited understanding of whether specific CAF rewiring is induced by different neoplastic cell states. We argued that the identification of a CAF transcriptional phenotype specifically associated with a biologically aggressive cancer subtype may improve patients’ stratification based on the risk, lead to a better prediction of therapeutic responses, and potentially guide effective personalized therapies. Activating mutations of *KRAS* are nearly universal in PDAC ^24^ and result in the hyperactivation of the RAS/MEK/ERK (Mitogen-activated protein kinase, MAPK) pathway, which plays a central role in tumour initiation and maintenance ^25,26^. Increased dosage of oncogenic *KRAS* is observed in basal-like/squamous subtypes ^17,27,28^, yet classical PDAC cells display superior sensitivity to the inhibition of MAPK ^29^. Targeting of MAPK, alone or in combination, has demonstrated to be ineffective in PDAC ^30,31^. However, the significance of MAPK activity in the PDAC tumour microenvironment has never been explored so far. Here, we uncovered an important role for the MAPK signalling pathway in the definition of PDAC CAF phenotypes. We show that the myCAF transcriptional phenotype is uniquely dependent on proficient MAPK signalling. Additionally, we demonstrate that hyperactivation of MAPK signalling occurs in myCAFs populating basal-like/squamous tumour niches with reduced frequency of CD8+ T cells. Gene expression signatures of MAPK^high^ CAFs from mouse tumours suggest immunoregulatory functions. Finally, we found that gene expression signature of MAPK^high^ CAFs correlates with poor prognosis in several human cancers, including PDAC, and with reduced response to immune checkpoint inhibition in bladder cancers and melanoma.

## RESULTS

### Basal-like PDAC cells bear cancer-associated fibroblasts with elevated MAPK activity

To identify molecular features predictive of cellular dependency on MAPK signalling, we treated 6 human cell lines and 5 patient-derived organoids (PDOs) with increasing doses of the MEK1/2 inhibitor trametinib (MEKi). In both cell lines and PDOs, the classical lineage and high levels of a transcriptional signature of MAPK activation (MAPK Biocarta) positively correlated with higher sensitivity to MEKi (**Supplementary Fig. 1a, b**). Regardless of the subtype, short-term treatment of PDAC cell lines with subIC50 doses of MEKi effectively inhibited MAPK (**Supplementary Fig. 1c**). Treatment of cell lines with sublethal doses of MEKi for 2 and 7 days did not induce changes in cell identity (**Supplementary Fig. 1d**). However, the basal-like cell line (Hs766t), which displays high sensitivity to MAPK inhibition, showed the largest change in transcriptome following treatment (**Supplementary Fig. 1e**). Then, we used the same transcriptional signature of MAPK activation to infer differential activity of the pathway in an extended set of human cell lines. We explored the transcriptomic data available from the Cancer Cell Line Encyclopedia (CCLE) (n = 41) ^32^ and those available in our laboratory (n = 10) to reliably classify cell lines as either basal-like or classical ^19^ (see methods, **Supplementary Table 1**). In this dataset, the MAPK transcriptional signature could not discriminate classical from basal-like cell lines (**Fig. 1a**). When exploring transcriptomic data from tissues, high levels of the MAPK signature were enriched in the basal-like PDAC of The Cancer Genome Atlas (TCGA)^16^ but not of the PanCuRx ^17^ or the International Cancer Genome Consortium (ICGC) ^15^ cohorts (Fig. 1a). To refine our analysis, we used RNA-seq data from 6 MEKi treated cell lines (Supplementary Fig. 1a) to derive a transcriptomic signature of MEKi response (epithelial MEK, eMEK) based on downregulated genes only, which we considered as a MAPK dependency footprint (**Supplementary Fig. 1f** and **Supplementary Table 2**). Consistent with our observations, the eMEK signature could not discriminate classical from basal-like tumours in the PanCuRx cohort ^17^, which contains RNA-seq data from microdissected epithelia (**Supplementary Fig. 1g**). The tissues included in the TCGA cohort display, on average, a very low neoplastic cell content ^16^. Therefore, we reasoned of a potential contribution of non-malignant cells to the elevated MAPK activation inferred in the basal-like tissues of this cohort. We focused on CAFs and tumour-associated macrophages as they represent the most abundant cell types in the PDAC TME. MAPK transcriptional signatures were positively correlated with the levels of stromal genes such as Podoplanin (*PDPN)*, Actin alpha 2 (*ACTA2),* and Fibroblast activation protein (*FAP)* (**Supplementary Fig. 1h**). Next, we applied a multiplex immunofluorescence (mIF) to human PDAC tissues classified as either basal-like/squamous or classical with well-established tissue markers ^15,33–36^ (**Supplementary Fig. 1i**, j). We found a significant enrichment for α-smooth muscle actin and phosphorylated extracellular signal-regulated kinase (α-SMA^+^p-ERK^+^) fibroblasts in the stroma of basal-like/squamous cells (**Fig. 1b**). In tumour tissues showing co-existence of classical and basal-like cells, we observed preserved connection between basal-like cells and p-ERK^+^ CAFs (**Fig. 1c**). In GATA Binding Protein 6 high (GATA6^high^) Keratin 81 negative (KRT81^neg^) (i.e., classical) tumour regions ^37^, p-ERK^+^ CAFs were rarely detected while the opposite was found in KRT81^high^-expressing tumour regions (Fig. 1c). Spatial analysis showed that significantly more α-SMA^+^p-ERK^+^ CAFs can be found in proximity of basal-like cells (Fig. 1c). Finally, we investigated if those p-ERK^+^ CAFs were anchored to the basal-like phenotypes using tissues from patient-derived xenografts (PDX) that exhibited class switch following drug treatment (n=5). We found that the treatment-induced switch of epithelial subtype from classical to basal-like was coherently associated with increased p-ERK signals in the stroma compartment (**Fig. 1d**). Conversely, no evident change was observed in the stromal compartment when there was no switch of tumour subtype upon treatment (n=10) (**Supplementary Fig. 1k**). Pancreatic stellate cells (PSCs) are known precursors of cancer-associated fibroblasts ^38–40^. We serum-starved mouse PSCs (mPSCs) and then briefly exposed them to the conditioned media from either classical or basal-like human PDAC cells before evaluating levels of p-ERK. The conditioned medium from basal-like cells induced nuclear enrichment of p-ERK to the same extent of mouse Epidermal growth factor (mEGF) and significantly more than the conditioned medium from classical PDAC cells (**Fig 1e**). Accordingly, the immunoblot analysis of mPSCs exposed to cell lines’ conditioned media confirmed enhanced p-ERK upon exposure to basal-like cancer cells secretome (**Fig. 1f**). Collectively, these data support a mechanistic link of basal-like PDAC cancer cells driving elevated activity of MAPK in CAFs.

**Figure 1.**
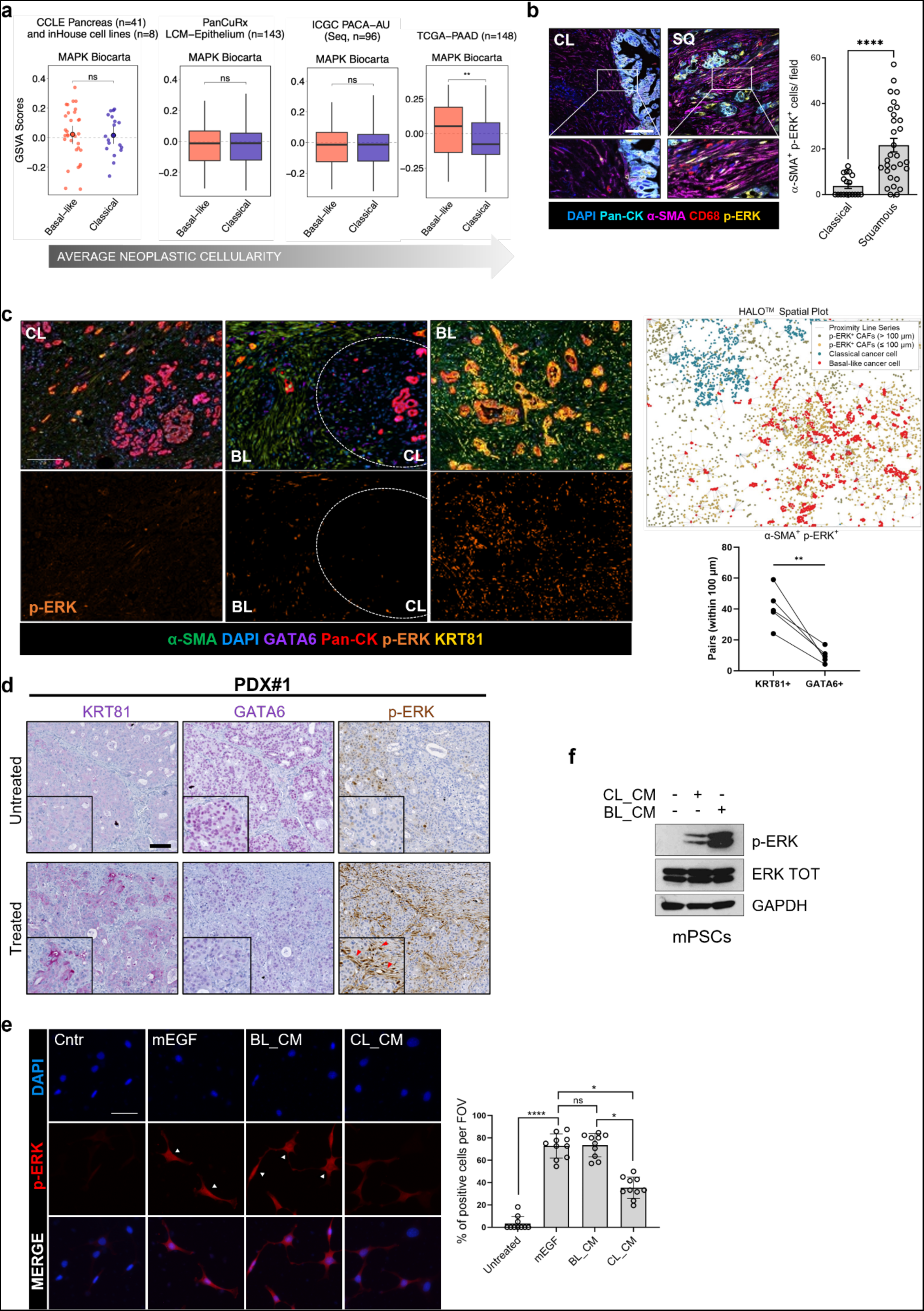
Basal-like PDAC cells bear cancer-associated fibroblasts with elevated MAPK activity. **a** Scatter dot plot and boxplots of GSVA scores obtained for the MAPK Biocarta gene set (MSigDB) stratified by Moffitt subtypes ^19^. From left to right: CCLE ^32^and in house cell lines, PanCuRx ^17^, ICGC ^15^, and TCGA ^16^. **p < 0.01; and ns, not significant as determined by Wilcoxon test. **b** Representative images of multiplex IF performed on FFPE of tumours of classical (CL) or squamous (SQ) subtype. Scale bar, 100 μm. Below, 2X magnification of the selected areas. The quantification is provided on the right as dot plot displaying the numbers of α-SMA^+^p-ERK^+^ cells per field of visualization (20X magnification areas). A minimum of 4 fields for case (5 cases/subtype) were analysed. ****p < 0.0001 as determined by Student’s t test (two-sided). Error bars show standard deviation. **c** Representative images of multiplex IF performed on FFPE of human PDAC tissues. The three panels represent different areas within the same tumour displaying different molecular phenotypes. CL, classical; BL, basal-like. Scale bar, 200 μm. On the right, the spatial plot obtained with Halo® Image Analysis Platform showing the distribution of p-ERK^+^ CAFs around classical (GATA6^+^) and basal-like (KRT81^+^) cancer cells. Below, the paired dot plot showing the quantification of p-ERK^+^ CAFs with a distance below 100 μm to classical and basal-like cancer cells (n=5). **p < 0.01 as determined by Student’s t test (two-sided). **d** Representative images of immunohistochemistry for KRT81, GATA6 and p-ERK on tissues from a patient derived xenograft (PDX#1) showing class switch upon MEKi treatment (n=4). Red arrows indicate p-ERK signal in stroma. Scale bar, 200 μm. Inserts showed 2X magnification of selected areas. **e** Representative immunofluorescence staining of p-ERK on mPSCs FBS-starved for 6 hours (Cntr) and treated for 10 minutes with mEGF (50 ng/ml) and basal-like or classical conditioned media. Nuclei were counterstained with DAPI (blue). Scale bar, 50 μm. White arrows indicate nuclear staining of p-ERK. On the right, scatter dot plot showing the percentage of cells with p-ERK nuclear staining per field of visualization (FOV; n=10/condition). ****p < 0.0001; *p<0.05, and ns, not significant as determined by one-way Anova. Error bars show standard deviation. **f** Immunoblot analyses of p-ERK and total ERK in whole cell lysates from mPSCs serum starved for 6 hours and treated for 10 minutes with either classical or basal-like conditioned media. GAPDH was used as loading control.

### scRNA-seq of mouse PDAC tumours treated with MEKi reveals quantitative and qualitative changes in cell subsets

To investigate the role of MAPK in the definition of stromal phenotypes, we performed a multidimensional analysis of tissues from mouse basal-like PDAC (**Fig. 2a**). First, we generated and characterized a mouse model based on the orthotopic transplantation of a quasi-mesenchymal KPC (Kras^G12D/+^; p53^R172H/+^; Pdx1-Cre) ^41^ derived cell line that *in vivo* produces cancer tissues aligning with the human basal-like PDAC phenotype ^42^. Tumour-bearing mice treated daily with 1 mg/kg of MEKi showed reduced activation of MAPK signalling (**Supplementary Fig. 2a**). Multiplex IF of tissues from tumour-bearing mice treated over the course of 14 days showed the presence of p-ERK^+^ CAFs in untreated tumours, the reduction of p-ERK signal in both the epithelial and stromal compartments at 2 days following treatment, and the rapid pathway rewiring particularly in the stromal compartment starting at 7 days of treatment (**Fig. 2b**). In keeping with that, the reactivation of the MAPK pathway occurred *in vitro* within hours from the treatment of mPSCs (**Supplementary Fig. 2b**). Given the kinetics of the MAPK pathway rewiring, we then performed scRNA-seq on fresh tissues from tumour-bearing mice treated for 2 and 7 days with MEKi along with their untreated controls. We profiled a total of 18,495 cells across 12 tumours (6 untreated and 6 treated, see methods) and recovered an overall cellular composition similar to that expected from PDAC tissues (**Fig. 2c, d**). Unsupervised clustering of single cell data identified 11 major cell types (Fig. 2c). Further annotation of the identified cell subsets was performed *post hoc* by using known gene signatures (Fig. 2d). The main cluster was almost exclusively composed by malignant epithelial cells which were confirmed by inferred copy-number alterations (CNA) ^43^ (**Supplementary Fig. 2c**). Other cell types included non-malignant epithelial cells (i.e., acinar cells), immune and stromal cell types (Fig. 2c, d). The treatment was associated with significant changes in cell composition and cell states. We observed a significant reduction in the proportion of malignant epithelial cells both at 2 and 7 days of treatment (**Fig. 2e, f**), which was consistent with the histology of treated tumours (**Supplementary Fig. 2d**). In the non-malignant compartment, we observed a significantly higher fraction of CAFs in the treated groups that was also confirmed with immunohistochemistry for the fibroblast marker FAP (**Supplementary Fig. 2e**). Consistent with the pharmacological treatment and proteomic analysis, the eMEK transcriptional signature was significantly downregulated in the epithelial compartment of treated tumours (**Fig. 2g**). MEKi was also associated with changes in the proportion of basal-like and classical neoplastic cells (**Fig. 2h** and **Supplementary Fig. 2f**). However, molecular subtyping on epithelial pseudo-bulk showed no significant changes in cell state, which is in line with the results obtained on treated human PDAC cell lines (Supplementary Fig. 1d).

**Figure 2:**
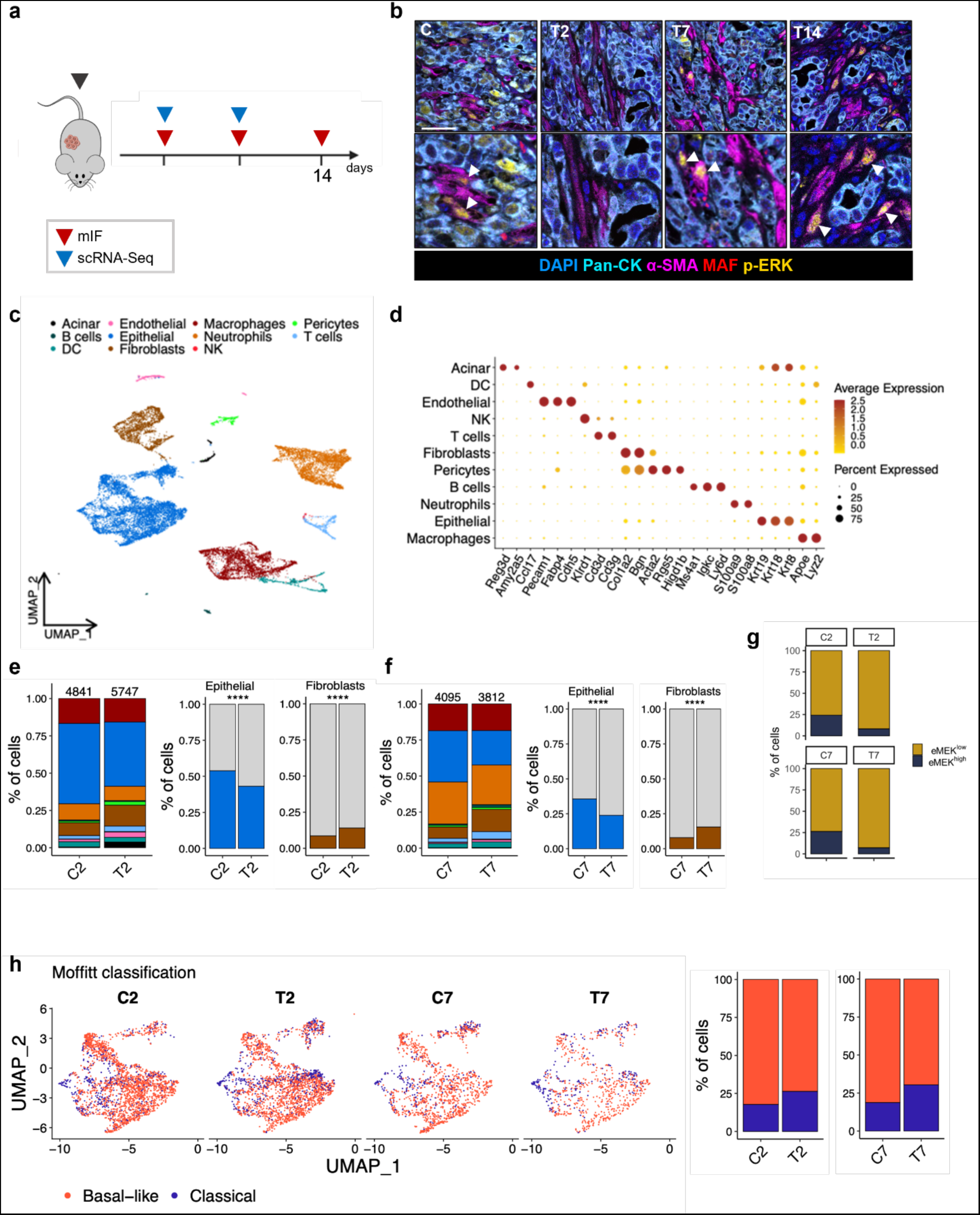
scRNA-seq of mouse PDAC tumours treated with MEKi reveals quantitative and qualitative changes in cell subsets. **a** Schematic representation of the *in vivo* experimental setting. At each timepoint, a group of mice have been sacrificed and further analysis performed. **b** Representative images of multiplex immunofluorescence performed on tumour tissues from mice treated with (from left to right): vehicle (C), MEKi for 2 (T2), 7 (T7), and 14 (T14) days. White arrowheads indicate α-SMA^+^/p-ERK^+^ cells. Bottom panels showed a 2X magnification of selected areas. Scale bar, 25 μm. **c** UMAP plot showing the unsupervised clustering of viable cells from 12 digested mouse tumours tissues (6 vehicle and 6 treated) annotated in 11 different clusters. Different cell type clusters are color-coded. **d** Bubble plot showing selected cell type-specific markers across clusters. Size of dots represents the percentage of cells expressing a specific marker and intensity of colour indicates level of average expression. **e-f** Barplots displaying the percentage of cells for each cell type cluster from mice treated with either vehicle (C) or MEKi (T) for 2 (e) or 7 (f) days. Numbers refer to the total number of cells for each sample. ****p < 0.0001 as determined by χ² test. **g** Barplot representing the percentage of cells of the epithelial cluster stratified for eMEK signature in vehicles and in MEKi treated samples. **h** UMAP plots showing cells of the epithelial cluster from vehicle (C2-C7) and MEKi (T2-T7) treated mice. Cells were classified as classical and basal-like according to Moffitt’s classification^19^. On the right, barplots showing the percentage of cells of the epithelial cluster of mice treated either with vehicle or MEKi for 2 or 7 days classified according to Moffitt’s classification ^19^.

### MAPK inhibition induces changes in the proportion of myCAFs and iCAFs

Established signatures of CAFs commonly found in mouse and human PDAC tissues ^6^ were expressed in distinct clusters of cells, which could be detected across conditions (treated/untreated) and timepoints (2 and 7 days) (**Fig. 3a**). The main subset of CAFs was composed by myCAFs, while iCAFs was the less represented one in all conditions (Fig. 3a). As observed for the malignant epithelial compartment, the treatment induced quantitative and qualitative changes in the fibroblast compartment. While the treatment did not partition CAFs into distinct clusters (**Supplementary Fig. 3a**), we observed a significant reduction in the proportion of myCAFs after treatment, which was associated with an increase in the number of iCAFs (**Fig. 3b**). These relative changes were more prominent at 2 days compared to 7 days following MEKi. Considering all CAFs subsets, the treatment induced the significant downregulation of the MAPK transcriptional signature (**Supplementary Fig. 3b).** However, the myCAFs compartment showed the largest variation in gene expression upon treatment, which suggests a higher dependency of this CAFs phenotype on MAPK activity (**Fig. 3c**). We next explored the possibility that MEKi could affect the differentiation status of CAFs. First, we used CytoTRACE ^44^ to predict the differentiation status of the two CAF subsets in untreated and treated tumours. Regardless of the treatment, iCAFs and myCAFs were at the opposite ends of the differentiation spectrum (**Fig. 3d**), with iCAFs showing the lowest transcriptional diversity and therefore predicted to be the more differentiated cell subset. Next, we used the velociraptor toolkit ^45^ to infer pseudotemporal trajectory from scRNA-seq data ^46^. In untreated tumours, no dominant pseudotrajectory could be identified (**Fig. 3e**). Conversely, a dominant pseudotrajectory from myCAFs to iCAFs was inferred in treated tumours, supporting MAPK as a relevant pathway for the maintenance of the myCAF phenotype (Fig. 3e). To orthogonally validate these findings, we first explored scRNA-seq from untreated tumours to identify reliable markers of myCAF and iCAF subsets. We found that *Tnc* and *Mmp3* were highly expressed in cell subsets showing myCAFs and iCAFs phenotypes, respectively (**Supplementary Fig. 3c, d**). *In situ* RNA hybridization analysis (ISH) of mouse PanIN from autochthonous KC (Kras^G12D/+^; Pdx1-Cre) models driven by oncogenic K-RAS ^47^ revealed expression of *Tnc* in proximity of epithelial cells while abundant stromal cells expressing *Mmp3* were found surrounding each lesion (**Supplementary Fig. 3e**). In cancer tissues from autochthonous KPC models driven by mutations of *K-ras* and *Trp53* ^41^, both *Tnc* and *Mmp3* expressing cells showed a spatial segregation consistent with that observed for myCAFs and iCAFs phenotypes in this model ^4,6,8^ (Supplementary Fig. 3e). Next, we performed RNA ISH in tissues from our mouse cohort. Consistent with the scRNA-seq data, the treatment induced statistically significant changes in the proportion of myCAFs and iCAFs (**Fig. 3f**). Specifically, after short-term MEKi, there was a drastic reduction in the number of *Tnc* expressing cells with a concomitant increase in *Mmp3* expressing cells (Fig. 3f). To further generalize our findings and understand whether the treatment-induced changes into myCAF to iCAF proportion occurred independently of the epithelial cell lineage, we analysed tissues obtained from syngrafts orthotopically transplanted from cell lines of classical subtype derived from *Kras^G12D^;Tp53^fl/fl^* background mice ^48^ (**Supplementary Fig. 3f, g**). Mouse PDAC tissues showed expression of the classical marker GATA6 and accordingly p-ERK staining was limited to the epithelial cells (Supplementary Fig. 3f). Short-term treatment with MEKi of tumour-bearing mice was consistently associated with a significant increase in iCAFs also in classical tumours (Supplementary Fig. 3g). Altogether, our data indicates that MAPK activity is a key determinant of the myCAF phenotype *in vivo*.

**Figure 3:**
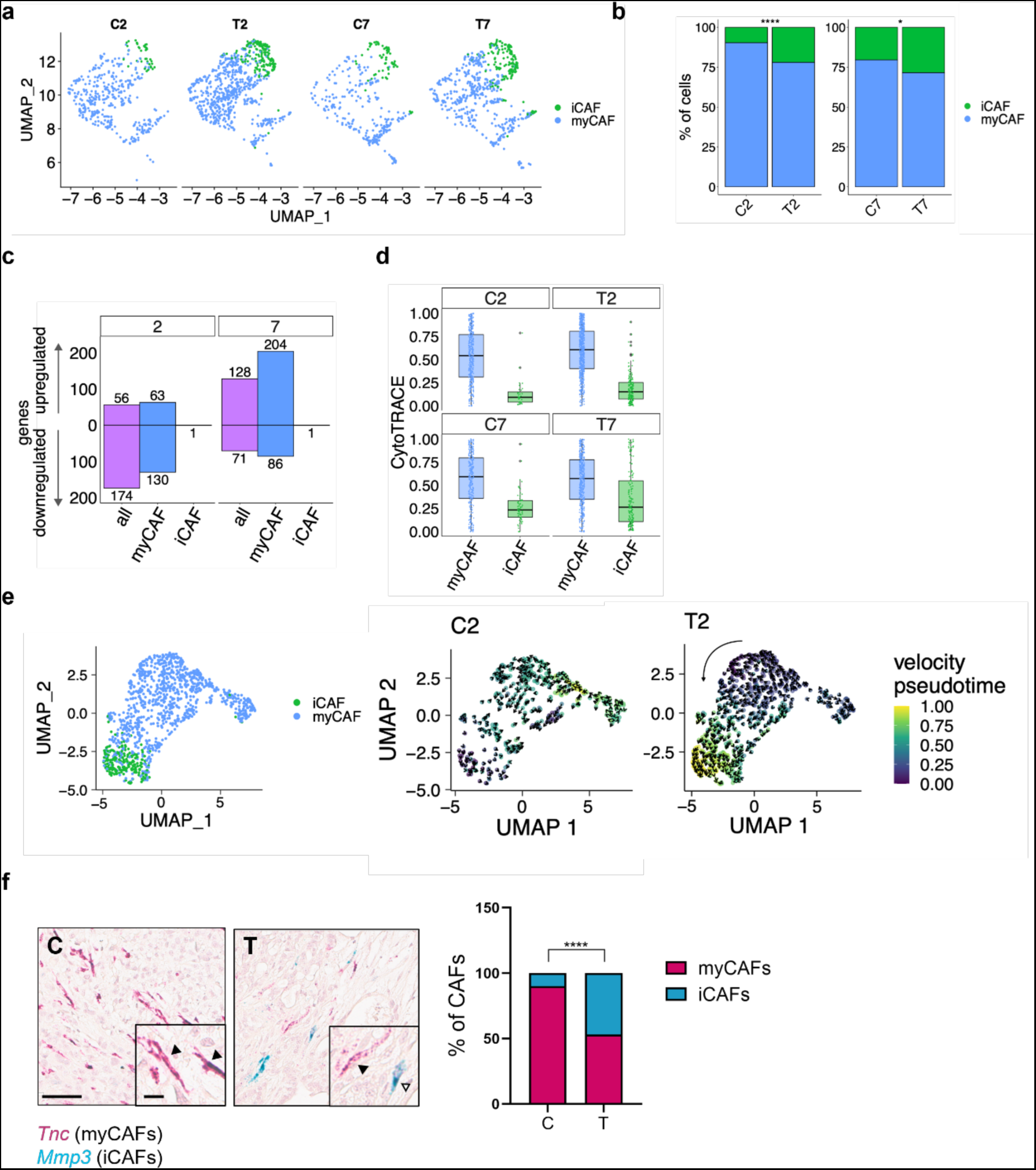
MAPK inhibition induces changes in the proportion of myCAFs and iCAFs. **a** UMAP plot of cells from the fibroblast cluster from mice treated with either vehicle or MEKi for 2 or 7 days. Cells were classified as either myCAFs or iCAFs according to Elyada’s classification ^6^. **b** Barplots representing the percentage of cells of the fibroblast cluster according to subtypes from Elyada et al. ^6^ ****p < 0.0001, *p < 0.05 as determined by χ² test. **c** Barplots showing the number of genes up- and down-regulated in cells of the fibroblast cluster after 2 and 7 days of treatment with MEKi. Cells are stratified using Elyada’s classification ^6^. **d** Scatterplot showing the CytoTRACE values ^44^ for cells of the fibroblast cluster treated either with vehicle or MEKi for 2 or 7 days and classified according to Elyada’s subtypes ^6^. **e** On the left, UMAP plot of the fibroblast cluster obtained by the integration of cells from mice treated with either vehicle or MEKi for 2 days. The colour visualization represents Elyada’s subtypes ^6^. On the right, UMAP representing the velocity (arrows) and pseudotime (colour) for each cell of the fibroblast cluster (annotated on the left panel) from mice treated with either vehicle (C2) or MEKi (T2) for 2 days. Black arrow indicated the global directionality of the velocity. **f** Representative images of *in situ* hybridization showing expression of *Tnc* (red; myCAFs) and *Mmp3* (green; iCAFs) genes on PDAC tissues from mice treated with either vehicle (C) or MEKi (T). Scale bar, 60 μm. Insert showed a 2X magnification of selected areas. Black arrowheads indicate myCAFs, while white arrowheads indicate iCAFs. Quantification is provided on the right as barplot displaying the percentage of iCAFs and myCAFs in tissues from mice (n = 5) treated with either vehicle or MEKi. ****p < 0.0001 as determined by Student’s t test (two-sided).

### A MAPK CAF signature identifies a subcluster of myCAFs and is associated to basal-like tumours

To identify subsets of CAFs with elevated activity of MAPK in our scRNA-seq data, we derived a transcriptomic signature based on genes significantly downregulated by two days of MEKi in the CAF compartment (stromal MEK, sMEK) (**Fig. 4a, Supplementary Fig. 4a**, and **Supplementary Table 3**). Next, we classified cells by quartiles of Gene Set Variation Analysis (GSVA) score distribution of the sMEK signature (n = 169 genes) and defined MAPK^high^ CAFs as cells in the highest quartile. This subset of CAFs formed a somewhat distinct cluster of cells in untreated tumours (**Fig. 4b**) and was exclusively composed by myCAFs (**Fig. 4c**). We then explored single cell RNA-seq data from cancer tissues of autochthonous KPC mice^6^ and consistently found that MAPK^high^ CAFs were mostly myCAFs (**Supplementary Fig. 4b**).

**Figure 4:**
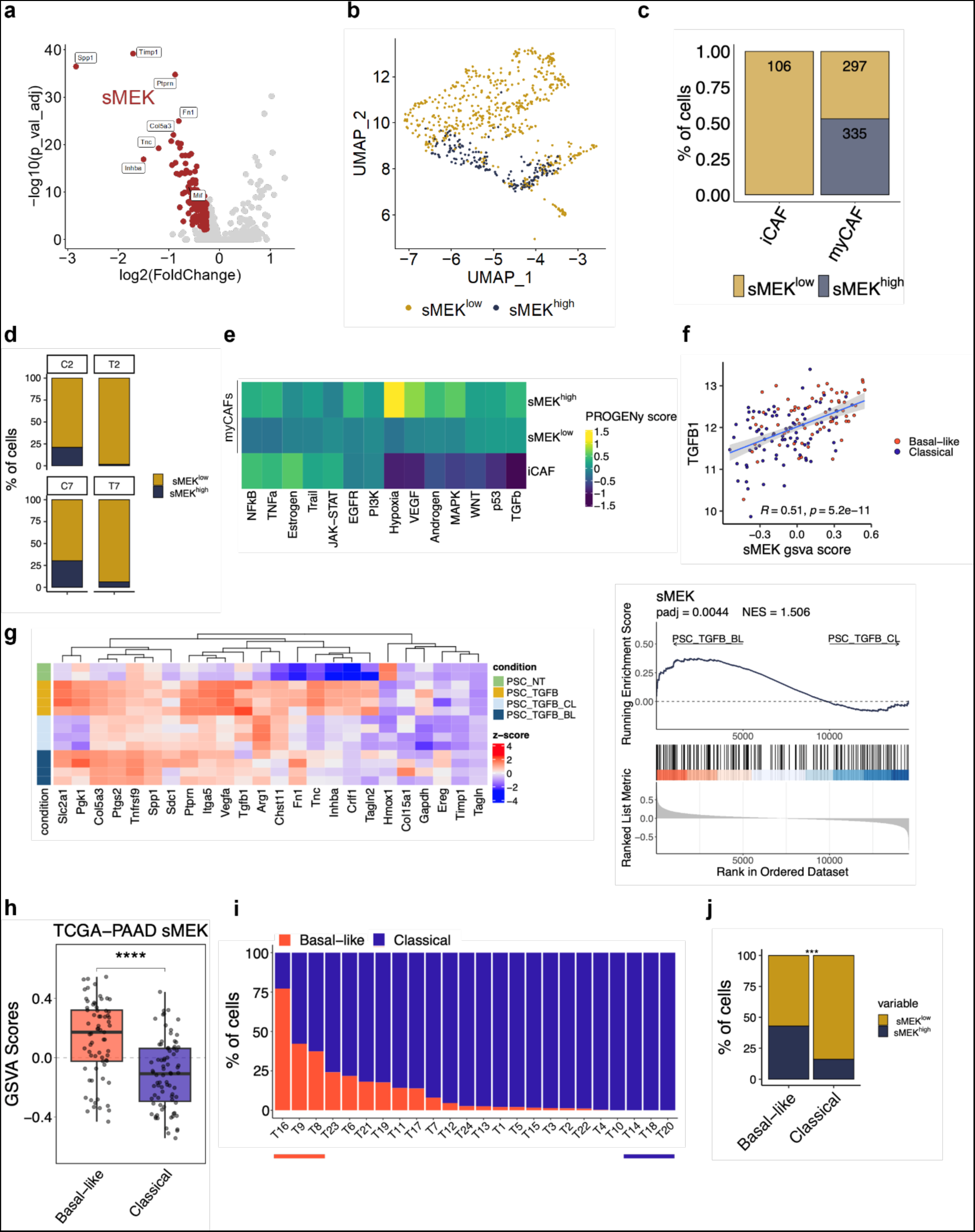
A MAPK^high^ signature identifies a subcluster of myCAFs enriched in basal-like tumours. **a** Volcano plot representing the differentially expressed genes in fibroblasts upon treatment with MEKi for 2 days. The red dots are the genes defining the sMEK signature (n=168). Highlighted some of the genes with log2FC expression < −2 and adjusted *p* < 0.05. **b** UMAP plot of cells from the fibroblast cluster stratified according to sMEK signature from mice treated with vehicle. **c** Barplot displaying the percentage of cells of the fibroblast cluster of untreated mice showing elevated MAPK activity (sMEK^high^) and classified as either myCAFs or iCAFs. **d** Barplot representing the percentage of cells of the fibroblast cluster stratified for sMEK signature in vehicles and in MEKi treated samples. **e** Heatmap showing expression of pathway-responsive genes in specific CAFs phenotypes as assessed by PROGENy analysis^44^. **f** Scatter plot showing the positive correlation between *TGFB1* expression and sMEK GSVA score for samples of the TCGA cohort ^16^. Samples were further stratified as either classical or basal-like subtypes from Moffitt et al. ^19^. **g** Heatmap showing changes in the expression of the genes of sMEK signature in mPSCs treated with either TGF-β1 (5 ng/ml) for 72 hours or pre-treated with TGF-β1 for 48 hours and then treated with conditioned media from classical or basal-like cell lines for 24 hours. On the right, GSEA plot showing the enrichment of sMEK signature in mPSCs treated with conditioned media from the basal-like cell line. **h** Boxplot of the GSVA scores for sMEK signature in samples of TCGA cohort ^16^ stratified by the Moffitt’s subtypes ^19^. ****p < 0.0001 as determined by Student’s t test (two-tailed). **i** Barplot showing percentages of cells of the epithelial cluster from Peng et al.^51^ classified as either classical or basal-like according to Moffitt’s subtypes ^19^. Cases are ranked by decreasing basal-like cells content. Samples displaying prevalent basal-like (n = 3) or classical (n = 3) epithelial cells were used in j. **j** Barplot showing the percentage of cells of the fibroblast cluster stratified according to the expression of sMEK signature in classical and basal-like samples from i. ***p < 0.001 as determined by χ² test.

Consistent with the dynamics of MAPK inhibition in our model (Fig. 2b), the proportion of CAFs expressing the sMEK signature decreased after 2 days of treatment and then increased at 7 days (**Fig. 4d**). Then, we used PROGENy ^49^ to infer pathway activity in each CAF subset from untreated tumours. In line with previous reports ^4–6,8,50^, myCAFs were characterised by elevated activation of TGF-β, while iCAFs were enriched for inflammation-associated signalling pathways (**Fig. 4e**). sMEK^high^ CAFs were characterised by elevated TGF-β and MAPK activity, the enrichment of hypoxia-driven responses, and activity of inflammatory pathways (Fig. 4e). Gene-set enrichment analysis (GSEA) of differentially expressed genes between CAFs with high and low MAPK activity confirmed metabolic reprogramming, as well as activation of inflammation-associated transcriptional programs in sMEK^high^ CAFs (**Supplementary Fig. 4c**). Accordingly, sMEK^high^ CAFs expressed high levels of Solute carrier family 2 member 1 (*Slc2a1),* Phosphoglycerate kinase 1 *(Pgk1),* Enolase 1 *(Eno1),* Pyruvate kinase M1/2 *(Pkm),* and some cytokine/chemokines, such as Interleukin 11 (*Il11)* and Macrophage migration inhibitory factor (*Mif)* (Supplementary Table 3). In addition, the sMEK signature positively correlated with *TGFB1* in the TCGA dataset ^16^(**Fig. 4f**). We next sought to understand whether similar transcriptional changes could be observed *in vitro*. mPSCs exposed to conditioned media from both basal-like and classical cancer cell lines showed induction of iCAF genes and activation of the JAK-STAT pathway, which was reported to be the driver of the iCAF phenotype ^4^ (**Supplementary Fig. 4d, e**). Therefore, mPSCs were pre-treated for two days with TGF-β1 to attenuate the polarization towards iCAFs and then exposed for 24 hours to either basal-like or classical CM. We observed that, along with stronger ERK phosphorylation, the basal-like CM induces the sMEK signature (**Fig. 4g**, Supplementary Fig. 4e). Of note, treatment with TGF-β1 upregulated the sMEK signature as well, in accordance with the myofibroblastic nature of sMEK^high^ CAFs (Fig. 4g). To translate our findings in the human setting, we explored bulk and scRNA-seq data from human PDAC tissues. The sMEK signature identified basal-like tumours from the TCGA cohort ^16^ (**Fig. 4h**) and, accordingly, was enriched in the stroma of human tumours displaying a greater proportion of basal-like cells (n=3) from the scRNA-seq dataset of Peng et al. ^51^ (**Fig. 4i, j**). In keeping with the mouse data, the sMEK^high^ signature was significantly enriched in the myCAF compartment of all tumours from Peng et al. ^51^ (**Supplementary Fig. 4f**).

### The sMEK signature can be found across several human cancer types and predicts poor clinical outcome

In the majority of the human PDAC cohorts investigated (n=4) ^52–55^, high levels of the sMEK signature outperformed established stromal signatures ^19^ in identifying patients with inferior overall survival (**Fig. 5a, b** and **Supplementary Fig. 5a**). Furthermore, we found that basal-like tumours displaying elevated levels of the sMEK signature had the worse prognosis (**Supplementary Fig. 5b**). Next, we performed a single-cell pan-cancer analysis across 7 tumour types ^56^ displaying myCAF and iCAF phenotypes in their TME ^57–63^. The sMEK signature was enriched in cancer-associated fibroblasts (i.e., activated fibroblasts) vs other cell types in most cancer indications (**Supplementary Fig. 5c**). Classifying the whole CAFs population of those tumours into either myCAFs or iCAFs, the sMEK signature was particularly prominent in the myCAF (**Fig. 5c**). In three tumour types (i.e., lung adenocarcinoma, bladder cancer, and uveal melanoma)^64–66^, elevated levels of the sMEK signature correlated with worse patient prognosis, as observed for PDAC (**Fig. 5d**). Notably, the sMEK signature was detected across several cancer types and correlates with *TGFβ1* expression (Supplementary Fig. 5c and Fig. 4f), a known predictor of poor response to immuno-oncology therapies ^67,68^. In addition to that, gene expression signatures of MAPK^high^ CAFs suggested immunoregulatory functions (Fig. 4a, Supplementary Fig. 4a, and Supplementary Table 3). Therefore, we first looked at density of cytotoxic T cells (CD8+) in PDAC tumour subdomains marked by p-ERK^+^CAFs. We found that p-ERK+ CAFs restricted the accumulation of cytotoxic T cells in their proximity (**Figure 5e, f** and **Supplementary Fig. 5d**). Next, we sought to test whether the presence of MAPK^high^ CAFs correlated with the response to cancer immunotherapy. The sMEK signature was enriched in patients with worse response to immunotherapy (stable and progressive disease) in patients with bladder cancer (BLCA) and malignant melanoma in the cohort from IMvigor210 trial^69^ and Hugo et al^70^, respectively (**Fig. 5g**). Accordingly, elevated level of the sMEK signature correlated with reduced survival in BLCA patients treated with anti–PD-L1 therapy ^5^ (**Supplementary Fig. 5e**). Overall, those data suggest that the sMEK signature is not PDAC-specific, correlates with poor outcome across different cancer types, and predict poor response to immunotherapy in immune-reactive tumours.

**Figure 5:**
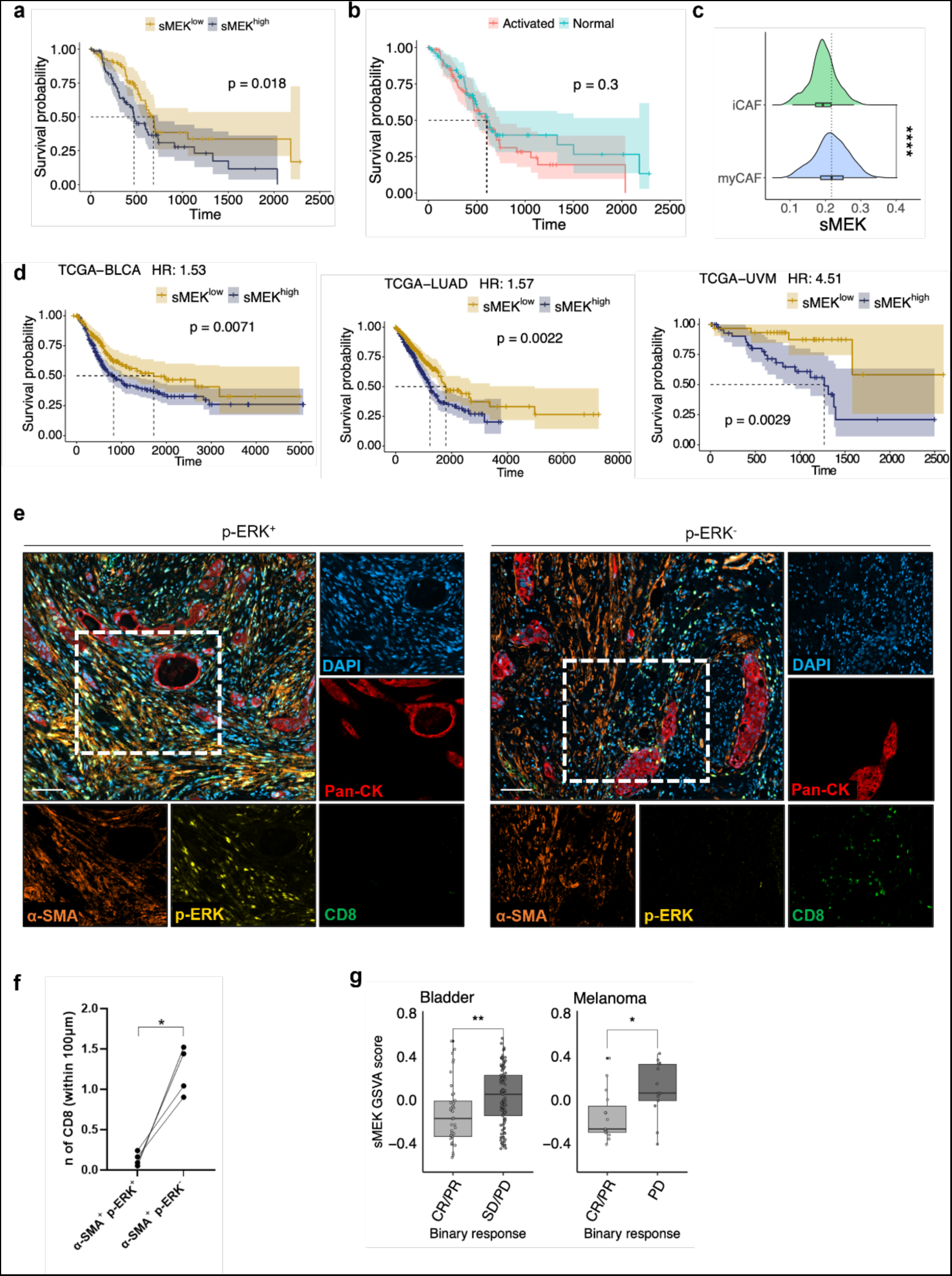
The sMEK signature predicts poor clinical outcome and resistance to immunotherapy. **a** Kaplan–Meier plot comparing the overall survival of patients from the TCGA cohort ^16^ according to the expression of sMEK signature. p, Log-rank (Mantel–Cox) test. **b** Kaplan–Meier plot comparing the overall survival of patients from the TCGA cohort^16^ according to the Moffitt’s stromal subtypes ^19^. p, Log-rank (Mantel–Cox) test. **c** Density plot for the sMEK signature score values over the activated fibroblasts from Luo et al. dataset ^56^. Activated fibroblasts were classified as either myCAFs or iCAFs. ****p < 0.0001 as determined by Student’s t test (two-tailed). **d** Kaplan–Meier plots comparing the overall survival of patients from bladder cancers^65^ (BLCA), lung adenocarcinomas ^64^ (LUAD), and melanoma^66^ (UVM) of the TCGA cohort according to the expression of sMEK signature. Displayed the annotated Cox hazard ratio (HR). p, Log-rank (Mantel–Cox) test. **e** Representative images of multiplex immunofluorescence of FFPE human PDAC tissue. The two panels show a p-ERK positive and a p-ERK negative area within the same tumour. Scale bar, 100 μm. **f** Paired dot plot showing the number of CD8^+^ cells with a distance below 100 μm to α-SMA p-ERK^+^ and α-SMA p-ERK^-^ CAFs. (n=4) *p < 0.05 as determined by Student’s t test (two-sided). **g** Boxplots showing sMEK GSVA score in bladder cancers from the IMvigor210 study ^69^ (left) and melanoma^70^ (right). Samples are separated by binary drug response. CR, complete response; PR, partial response; SD, stable disease; PD, progressive disease. ** p < 0.01, * p < 0.05 as determined by Wilcoxon test.

## DISCUSSION

Here, we undisclosed an important role for the MAPK signalling pathway in the definition of PDAC CAF phenotypes. Our data show that the myCAF transcriptional phenotype is uniquely dependent on a proficient MAPK signalling. Furthermore, hyperactivation of MAPK signalling occurs in myCAFs populating basal-like/squamous tumour niches which show reduced CD8+ T cells density. Differentiating the dependency of malignant and non-malignant cell populations on MAPK activity in PDAC is critical considering the recent advent of direct K-RAS inhibitors which specifically target mutant K-RAS driven MAPK activity^71^.

Gene expression signatures of MAPK^high^ CAFs from mouse tumours suggested metabolic rewiring and immunoregulatory function. We found that the sMEK signature correlates with poor prognosis in several cancers, including PDAC, and with reduced response to immune checkpoint inhibition in bladder cancers and melanoma. Our findings support previous observation about the importance of MAPK signalling in sustaining the myCAFs phenotype ^72^ and expand on the heterogeneous phenotypes that can be found in the PDAC TME. We provide new insights on a stromal phenotype specifically shaped by basal-like/squamous cells. Signalling pathway activities and cell dependency on a given pathway are often successfully inferred from gene expression data ^49,73^. Here, we used pathway mapping analysis and context-dependent pathway response signatures to infer MAPK activity and dependency in heterogeneous expression data from models and patients’ samples. In both human cancer cell lines and patient-derived organoids, high levels of a MAPK transcriptional signature were the best predictor of sensitivity to MEK1/2 inhibition. Basal-like/squamous models were predicted to be less dependent on MAPK activity and accordingly displayed reduced sensitivity to MEKi. *In vitro*, a transiently disabled MAPK pathway was not associated with significant changes of cancer cell states. Our results are in line with previous observation from Miyabayashi and colleagues showing that RAS signalling hyperactivation is not fundamental for the definition of the basal-like/squamous subtype in the epithelial compartment ^27^. We have previously shown that querying bulk transcriptional datasets from tissues with different neoplastic cell content is a viable strategy to localise the cellular compartment contributing to gene expression differences between molecular subtypes ^74^. Using the same approach, we show that MAPK transcriptional signatures were particularly elevated in basal-like tumours from the TCGA cohort^16^. The inferred hyperactivation of MAPK in the stromal compartment of tumours classified as basal-like significantly correlated with increased density of fibroblasts displaying nuclear p-ERK, a known surrogate marker for MAPK activation^75^.

Different neoplastic cell states often co-exist in human PDAC tissues ^17,21,76,77^. Therefore, molecular subtyping based on bulk sequencing data might mask transcriptional heterogeneity ^17,22^. Immunophenotyping of heterogeneous human tumours clearly showed that p-ERK^+^ CAFs were significantly enriched in basal-like/squamous niches. These p-ERK^+^ CAFs can be detected in mouse PDAC tumours as well as in heterospecies models, which were instrumental to demonstrate that this CAF subpopulation is specifically associated to the basal-like subtype. Indeed, treatment induced changes in cancer cell lineage (i.e, classical to basal-like) was accordingly associated with qualitative changes in the stroma, i.e., increased, or decreased p-ERK^+^ CAF density. To gain further insights into this CAF phenotype, we leveraged scRNA-seq data obtained from mouse tumour tissues following short-term perturbation of the MAPK pathway. Our model aligned with the human basal-like/squamous PDAC ^42^ and showed rapid kinetics of pathway rewiring in the stromal compartment following treatment. Short-term perturbations are often used to capture primary transcriptional response to a specific stimulus ^49^. Contrasting the two conditions with known differential pathway activity, we were able to show the general dependency of myCAFs on MAPK activity and to obtain gene-expression signatures of CAFs displaying MAPK hyperactivation. Our model preserved the CAF heterogeneity reported in mouse and human tumours ^4,6,8^, the expected myCAF/iCAF ratio in the PDAC TME ^1,5^, and showed presence of p-ERK^+^ CAFs, as expected for a basal-like model. Inferred dynamics in scRNA-seq data and RNA *in situ* expression analyses showed that MAPK inhibition leads to dramatic changes of the myCAF/iCAF ratio into the mouse PDAC TME. The highest degree of dependency of myCAFs on MAPK activity suggests a potential transdifferentiation of myCAFs into iCAFs. However, we cannot exclude those relative changes into fibroblasts composition in treated mice were also contributed by qualitative changes into the malignant compartment. Focusing on CAFs showing the highest dependency on MAPK activity, we obtained a gene expression signature (sMEK) that mapped almost exclusively onto the myCAF subcluster. This result is coherent with the spatial localization of p-ERK^+^ CAFs in human tissues and with previous evidence that locate myCAFs in close proximity to cancer cells^8^. Accordingly, the sMEK signature is enriched for ECM related genes and TGF-β driven programs. Nonetheless, the sMEK signature also suggested a glycolytic phenotype and immunoregulatory functions. Aligning with their myCAF identity, TGF-β signalling was a major driver of MAPK^high^ CAFs and accordingly treatment of mouse PSCs with TGF-β1 greatly induced the sMEK signature *in vitro*. Furthermore, the signature was also strongly induced when mPSCs were exposed to the conditioned media from basal-like human cancer cell lines. That further suggests that this phenotype is anchored to basal-like cancer cells. In different PDAC cohorts, the sMEK signature reliably identified patients with worse prognosis. That is in contrast with the epithelial specific MEK signature which failed to stratify patients based on risk. Previous studies have shown that similar fibroblast lineages and phenotypes can be observed in different cancers ^1,60,78^. We explored pan-cancer scRNA-seq data ^56^ to find that the sMEK signature is enriched in stromal cells from many cancer conditions and identifies more aggressive disease in some of them^64–66^. While deeper mechanistically investigations are needed, our data suggest potential immunoregulatory functions for MAPK^high^ CAFs. In keeping with that, p-ERK^+^ CAFs demarcated tumour areas showing reduced density of CD8+ T cells. The sMEK signature includes genes with immune-related function and high levels of the signature predict poor response to immunotherapy in bladder cancers and melanoma. In sum, our study shows that MAPK signalling is a key determinant of the myCAF phenotype and that its hyperactivation is a distinctive feature of stromal cells in basal-like/squamous tumour niches. Inhibition of MAPK signalling using a potent MEK1/2 inhibitor has important consequences on stroma remodelling with transient changes in myCAFs to iCAFs ratio. We also provide here a gene-expression signature that might be used for patients’ stratification based on risk in PDAC and other malignancies.

## MATERIALS AND METHODS

### Human samples

Human PDAC tissues used in this study were obtained from surgical resections of patients treated at the University and Hospital Trust of Verona (Azienda Ospedaliera Universitaria Integrata, AOUI). Written informed consent was acquired from patients before specimens’ acquisition. The FFPE samples used for staining were retrieved from the ARC-Net Biobank and were collected under the protocol number 1885 approved by the local Ethics Committee (Comitato Etico Azienda Ospedaliera Universitaria Integrata) to A.S. (Prot. 52070, Prog. 1885). Tissues from surgical resection used for the generation of organoids were collected under the protocol number 1911 approved by the local Ethics Committee (Comitato Etico Azienda Ospedaliera Universitaria Integrata) to V.C. (Prot. n 61413, Prog 1911 on 19/09/2018). All experiments were conducted in accordance with relevant guidelines and regulations. The Essen cohort is a retrospective study carried out according to the recommendations of the local ethics committee of the Medical Faculty of the University of Duisburg-Essen. Clinical data were obtained from archives and electronic health records. Patients who had undergone pancreatic resection with a final histopathologic diagnosis of human PDAC between March 2006 and February 2016 was used (Approval no: 17-7340-BO).

### Cell lines and organoids

We used 5 mouse PDAC cell lines, 1 mouse pancreatic stellate cell line, 11 human PDAC cell lines, and 5 human PDAC organoids. The mouse PDAC cell line FC1199 was generated from tumour of *KPC* mice (Kras^G12D/+^; p53^R172H/+^; Pdx1-Cre) ^41^. FC1199 were provided by the Tuveson’s laboratory (Cold Spring Harbor Laboratory, NY, USA) and was cultured in DMEM supplemented with 10% fetal bovine serum (FBS) and 1% Penicillin-Streptomycin (Pen-Strep). Primary murine PDAC cell lines 60400 and 110299 were derived from corresponding tumour pieces of *CKP* mice (60400: Ptf1a^wt/Cre^;Kras^wt/LSL-G12D^;p53^fl/fl^ ^79^; 110299: Ptf1a^wt/Cre^;Kras^wt/LSL-G12D^;p53^LSL-R172H/fl^ ^80^) and were cultured in DMEM high-glucose medium supplemented with 10% FBS. The mouse pancreatic stellate cell line (mPSC4) has been established from WT C57BL/6J mice ^8^. mPSC4 was provided by the Tuveson’s laboratory (Cold Spring Harbor Laboratory, NY, USA) and cultivated as previously described^8^. Human PDAC cell lines (PANC-1, HPAF-II, hT1, hM1, hF2, and Hs766T), human primary PDAC monolayer cell lines (VR2-2D (PDA2-2D), VR6-2D (PDA6-2D), VR9-2D (PDA9-2D), VR20-2D (PDA20-2D), and VR23-2D (PDA23-2D)) and human PDAC organoids (hT3, VR1-O (PDA1-O), VR2-O (PDA2-O), VR9-O (PDA9-O), and VR20-O (PDA20-O)) were obtained and cultivated as previously described^74^. Cell lines and organoids were routinely screened for Mycoplasma contamination using MycoAlert Mycoplasma Detection Kit (Lonza).

### Generation of mouse models

In this study we used both isograft and xenograft models. Six- to eight-weeks old C57Bl/6J (B6J) and NSG (NOD.Cg-Prkdc^scid^;Il2rg^tm1Wjl^) mice were purchased from Charles River Laboratory (Milan). All animal experiments regarding transplanted mice were conducted in accordance with procedures approved by CIRSAL at University of Verona (approved project 655/2017-PR). KC (Kras^G12D/+^; Pdx1-Cre) and KPC (Kras^G12D/+^; p53^R172H/+^; Pdx1-Cre) mice were used as spontaneous model for pre-invasive lesions and PDAC, respectively ^41,47^. Isograft models were generated with KPC and CKP-derived cell lines. For the generation of isograft based on KPC-derived cell line (FC1199), 2.5×10^5^ cells were transplanted in B6J mice as previously described^42^. For the generation of isograft based on CKP-derived cell lines (60400 and 110299), 5.0×10^5^ cells were resuspended in 30 µL of a 1:1 dilution of Matrigel® (Corning) and cold plain medium and injected into the pancreas of B6J mice using insulin syringes (BD micro-fine 30 Gauge) under the guidance of Ultrasonic imaging. The injection was considered successful by the appearance of encapsulated cell suspension ball without signs of leakage. Patient-derived xenografts (PDX#1, PDX#2) were generated by subcutaneous implantation of a patient’s tumour fragment in the left flank of anaesthetized NSG mice. Tumour growth was measured twice weekly until they reached the volume of 1 cm³. Then, tumours were harvested, cut into small fragments (3 mm^3^), and transplanted subcutaneously into the left flank of anaesthetized NSG to generate the 2^nd^ generation of xenograft. The same procedure was followed to generate the 3^rd^ generation which was used for treatments. Mice were maintained under sterile and controlled conditions (22°C, 50% relative humidity, 12 h light–dark cycle, autoclaved food and bedding, acidified drinking water).

### *In vivo* drug treatments

Isografts and xenografts were treated with Trametinib (MEKi) as indicated. Only female mice were used for treatments. Before treatment, tumour masses were measured, and mice were randomized. Xenografts and KPC-derived isografts were treated with Trametinib dissolved in a solution of 0.5% hydroxypropyl-methylcellulose, 0.2% tween 80 and ddH2O (pH 8) with a final concentration of 1 mg/kg for daily oral administration. CKP-derived isografts were treated with Trametinib dissolved in a solution of 1% Kolliphor® EL, 1% PEG400 and ddH2O with a final concentration of 1mg/kg for daily oral administration. Monitoring of tumour growth was performed as previously described ^81^. Xenografts were sacrificed after 4 weeks of treatment and isografts were sacrificed at the indicated time points. Pancreas was collected for downstream analysis.

### *In vitro* drug treatments

Cells and organoids were treated with Trametinib as indicated. Trametinib (Selleck) was dissolved in DMSO, whose final concentration was less than 0.1% (v/v). For each cell culture, the IC50 concentration was determined by the luminescence ATP-based assay CellTiter-Glo (Promega, G9683) following the manufacturer’s instructions. Briefly, 1×10^3^ cells were plated on white 96-well plate in 100 µL of culture medium. After 24 hours, cells were treated with increasing doses of Trametinib, and viability was measured at endpoint (72 hours) using a microplate reader (BioTek, Synergy 2 Multi-mode Microplate Reader). Organoids were dissociated in single cells as previously reported ^82^ and 1×10^3^ cells were plated on white 96-well plate in 20 µL of a 1:1 solution of Matrigel® (Corning) and splitting medium (Advanced DMEM/F12 medium supplemented with HEPES, Glutamax^TM^ and Primocin (1 mg/mL)). 80 μl of human complete medium ^83^ supplemented with Rho Kinase Inhibitor (10.5uM, RhoKi) were added in the well. After 36 hours, cells were treated with increasing doses of Trametinib, and viability was measured at endpoint (72 hours) using a microplate reader (BioTek, Synergy 2 Multi-mode Microplate Reader). For measurement of signalling pathways following pharmacological treatment, individual cell lines were counted and seeded in a 6 well plate or a 100 cm^2^ dish. After reaching 40% of the confluence, cells were treated for the time reported in each experiment. PDAC cell lines were treated with subIC50 concentration: HPAF-II (IC50 = 4.8 nM), PANC-1 (IC50 = 10000 nM), Hs766t (IC50 = 2.85 nM), hF2 (IC50 = 4.54 nM), hM1 (IC50 = 1.35 nM), hT1 (IC50 = 148.8 nM). mPSC4 have been treated with MEKi with a subIC50 concentration of 10 nM.

### Conditioned media treatments

Conditioned media was collected from cancer cell lines after 48 hours of conditioning and used to treat mPSCs as indicated in figure legends. Only when FBS depletion was required, mPSCs were treated with media collected from cancer cell lines after 15 hours of conditioning in serum starvation.

### Immunohistochemistry

The following primary antibodies were used on mouse PDAC tissues: p-ERK (#9101; #4376, Cell Signaling Technology), GATA6 (#AF1700, Bio-techne; #ab175349, Abcam) KRT81 (sc-100929, Santa Cruz Biotechnology Inc.), and FAP (#ab53066, Abcam). The following primary antibodies were used on human PDAC tissues: PDX1 (#ab134150, Abcam), CK5 (#XM26, Novocastra), ΔNP63 (ACI3066C, Biocare), S100A2 (#109494, Abcam), GATA6 (#AF1700, Bio-techne), and p-ERK (#9101, Cell Signaling Technology). Quantification of FAP was performed in at least ten random non-overlapping fields of visualization (FOV, 20x) in each tissue using ImageJ. To measure the percentage of brown pixels, DAB^+^ particles were counted automatically by the software after colour deconvolution of the image. The percentage is relative to the number of nuclei present in each of the selected areas.

### Multiplex Immunofluorescence

Opal Multiplex IF Kit (Akoya) was used to perform multiplex immunofluorescence according to manufacturer’s instructions. The following antibodies have been used on human PDAC tissue of figure 1b or mouse PDAC tissue from figure 2b: pan-Keratin (#4279, Cell Signaling Technology), p-ERK (#9101, Cell Signaling Technology), α-SMA (#ab5694, Abcam), CD68 (#76437, Cell Signaling Technology), and MAF (#A300-613A, Bethyl). Images were acquired with Leica TCS SP5 laser scanning confocal (Leica) and digitalized by the Leica Application Suite X (LAS X) software. Quantification of α-SMA^+^p-ERK^+^ cells was performed in a minimum of 4 random non-overlapping FOV (80x) for case (5 cases/subtype) by counting the number nuclei/cells with staining positivity for each selected area. Opal Multiplex system (Perkin Elmer, MA) was used to perform multiplex immunofluorescence according to manufacturer’s instructions. The following antibodies have been used on human PDAC tissues of figures 1c, 5e and supplementary figure 3f: pan-Keratin (#ab6401, Abcam), p-ERK (#4376, Cell Signaling Technology), α-SMA (#ab5694, Abcam), GATA6 (#ab175349, Abcam), KRT81 (#sc-100929, Santa Cruz Biotechnology), and CD8 (#ab101500, Abcam). Slides were scanned and digitalized by Zeiss Axio Scanner Z.1 (Carl Zeiss AG, Germany). Quantification of individual and/or co-expressing markers, and spatial interaction analysis in the mIF images was performed with HALO (Indica Labs).

### Spatial imaging analysis

The binary information of cellular and nuclear signals was co-registered after generating intensity thresholds for each fluorescent channel. Java and R algorithms were used to automatically analyse cell distances. Using ImageJ, overlapping mask regions were employed to identify cells, which were marked with a point at the centre of the DAPI+ cell nucleus. This analysis was performed by using the R Package Spatstat1 for converting cell coordinates into Euclidian distances between cells. Using the algorithm, any two cells are ranked according to their average distance and number of unique neighbours. The threshold to pair up two cells was set to 100µm.

### Immunofluorescence on mPSCs

5×10^3^ mPSCs were plated in a cell culture chamber slide. After 48 hours, cells were serum starved for 6 hours and treated for 10 minutes with conditioned media from FBS-starved cancer cell lines. mEGF (50 ng/ml) was used as positive control for p-ERK nuclear translocation. After treatment, the IF was performed following the suggested protocol for p-ERK antibody (#9101, Cell Signaling Technology). Quantification was performed by counting cells with nuclear signal for p-ERK per FOV divided by the total number of cells counted in the field (10 FOV/condition).

### In Situ Hybridization

The *in-situ* hybridization (ISH) was performed on 4 μm section of mouse tissues. Briefly, slides were deparaffinized in xylene for 10 minutes followed by 100% ethanol for 2 minutes. After drying, slides were first incubated for 10 minutes with RNAscope® Hydrogen Peroxide (Advanced Cell Diagnostics) and then for 15 minutes at 99°C with RNAscope® 1X Retrieval Reagents (Advanced Cell Diagnostics). After dehydration in 100% ethanol, slides were dried and incubated at 40°C for 20 minutes with RNAscope® Protease Plus (Advanced Cell Diagnostics). The RNAscope® Probes (Mm-Mmp3 and Mm-Tnc-C2, Advanced Cell Diagnostics) were added to the slides following RNAscope® 2.5 Duplex Detection Reagents kit’s instructions. Positive control probe 2.5 Duplex Positive Control Probe-Mm and 2-plex Negative Control Probe (Advanced Cell Diagnostics) were used as positive and negative control, respectively. Quantification was performed by counting the total number of spindle-like cells with signal for *Tnc* or *Mmp3* in each tumour section.

### Immunoblotting

Protein lysates were obtained from cells using Lysis Buffer (Cell Signaling Technology) supplemented with phosphatases and proteases inhibitors (PhosSTOP™ and cOmplete (TM) Mini Protease Inhibitor Co, Roche). After electrophoretic separation, proteins were transferred on a PVDF membrane and incubated with the following antibodies: p-ERK (#9101, Cell Signaling Technology), total ERK (#9102, Cell Signaling Technology), p-AKT (#4060, Cell Signaling Technology), total AKT (#9272, Cell Signaling Technology), p-S6 (#D57.2.2E/4858, Cell Signaling Technology), total S6 (#54D2, Cell Signaling Technology), p-STAT3 (#9145, Cell Signaling Technology), and total STAT3 (#12640, Cell Signaling Technology). Vinculin (#4650, Cell Signaling Technology) and GADPH (#5174, Cell Signaling Technology) were used as loading controls.

### RNA sequencing and data processing

RNA was extracted from cell lines and organoids using TRIzol (Life Technologies), followed by column-based purification with the PureLink RNA Mini Kit (Ambion). The quality of purified RNA samples was determined using a Bioanalyzer 2100 (Agilent) with RNA 6000 Nano Kit. RNAs with RNA Integrity Number (RIN) values greater than 7.5 were used to generate sequencing libraries using the TruSeq Stranded Total RNA Kit (Illumina) following manufacturer’s instructions. Libraries were prepared from TrueSeq Stranded RNA Kit (Illumina) and sequenced on Illumina instruments. After quality control and adaptor trimming, PSC reads were aligned to the GRCm38 genome using Salmon v1.4.0 ^84^. O^85^rganoids and cell lines reads were aligned to the GRCh38 genome using STAR v2.7^86^. Transcripts’ quantification was imported in R through tximport package v4.0 and raw counts were normalized using the R/Biocondutor package DESeq2 v1.30.0 ^85^. Differentially expression analysis has been performed using DESeq2 ^85^. The eMEK signature was derived by filtering the results of differential gene expression performed with DESeq2 ^85^ between cell lines untreated and treated with MEKi for 2 days, filtering according to pvalue<0.01 and log2FoldChange<-1. Only protein coding genes defined in EnsDb.Hsapiens.v86 package annotation were retained ^87^. GSVA R package v1.38.2 ^88^ was used to calculate the main PDAC transcriptomics subtypes gene set scores ^19^.

### Statistical Analysis and Data mining

Seven transcriptomic datasets from either cell lines or cancer tissues were used for data mining ^15,16,64,66,17,32,65^. Samples belonging to the ICGC ^15^ and the TCGA-PAAD datasets ^16^ were restricted to 69 and 148, respectively, by selecting only true PDAC cases. For survival analysis, four additional PDAC datasets were included comprising either RNA-seq ^53^ or microarray data^52,54,55^. All RNA-seq raw count data were managed and normalized with DESeq package ^85^. Affymetrix (microarray) data were managed and normalized using the oligo ^89^ and affy ^90^ packages. Survival analysis has been performed through the R packages survival v3.5.0 (https://CRAN.R-project.org/package=survival) and the graphical representation done with survminer v0.4.9 ^91^. GraphPad Prism was used for graphical representation of data. Statistical tests were performed with R or GraphPad Prism and are reported in each figure legend.

### Mouse single cell RNA sequencing

#### Sample preparation and sequencing

Single cell RNA sequencing was performed on digested PDAC tissues from mice treated with either vehicle or MEKi for 2 or 7 days. For the digestion, tumour samples were collected in Splitting Medium (AdDMEM/F12 medium supplemented with HEPES (10 mM), Glutamax, and Pen/Strep) supplemented with 0.1% BSA and RhoKi (10.5 uM). After washing with PBS, specimens were cut in small pieces (1mm^3^) and incubated for 20 minutes in a tube rotator at 37°C in warm Digestion Medium (PBS 1X, 2mg/mL Dispase I, 1.25mg/mL Collagenase Type II, 100ug/mL DNAse I, and 0,05% FBS) supplemented with RhoKi. The cell suspension was pipetted and incubated on ice to let the larger tissue clumps settle to the bottom of the tube. The surnatant was collected, spun down and the pellet was resuspended in Splitting Medium supplemented with 0.1% BSA and 10 mg/ml Soybean trypsin inhibitor and stored on ice (Fraction 1). Then, the larger undigested clumps were digested again for 10 minutes, and every step previously described was repeated twice until the collection of Fraction 2 and 3. After digestion, Fractions 1, 2, and 3 were combined, filtered (40um nylon cell strainer) and centrifuged. The pellet was then resuspended in ACK lysing buffer supplemented with DNAse I to remove red blood cells from the sample and spun down. Cells were washed with PBS supplemented with 10% FBS and 1×10^4^ cells (concentration 1.000 cells/ul) were submitted for sequencing. To generate single cell GEMs, cellular suspensions from 3 mice/condition were loaded on a GemCode Single Cell Instrument (10x Chromium System) and libraries were generated with GemCode Single Cell 3’ Gel Bead and Library Kit v3 (10x Genomics). After barcoding, GEMs were broken, and cDNA was cleaned with with DynaBeads MyOne Silane Beads. cDNA was amplified, cleaned with the AMPure beads and the quality was checked using Fragment Analyzer HS NGS Assay. Libraries were quantified by quantitative PCR (qPCR) (KAPA Biosystems Library Quantification Kit for Illumina platforms) and the sequencing was performed on NextSeq500 (Illumina) with 75 paired-end kit.

#### Data processing

Binary base call (BCL) files were processed with the 10X proprietary software Cell Ranger ^92^, with default and recommended parameters. FASTQs files were aligned to reference transcriptome GRCm38 by count pipeline. Counts matrices for all samples were imported with Seurat ^93^. Cells with low quality were filtered from counts matrices (200 <n° of genes x cell< 9000 & %mitochondrial gene count < 25%). Vehicle and treatment datasets were integrated using Seurat integration pipeline ^93^, clustering analysis were run on integrated dataset with Seurat FindCluster ^93^ function using a resolution of 1. Annotation of the dataset were performed looking at the expression of well-known cell type markers. Copy number analysis was performed on epithelial compartment with <u>In</u>ferCNV R package ^94^ using as reference the non-epithelial cells.

#### Subtyping and enrichment analysis

Subtyping of epithelial cells was performed using signatures from Moffitt et al.^19^, Bailey et al.^15^, and Collisson et al.^15^. Subtyping of fibroblasts was performed using signatures from Elyada et al.^6^. To assess cells subtype, an enrichment score was assigned for each gene set to each cell using UCell R package ^95^, the cell subtype was assigned based on maximum score achieved by a cell for a specific gene set. Other tested signatures were obtained through the msigdb packages ^96,97^. The related single cell enrichment score was computed with UCell package^95^.

#### Analysis of the fibroblast subcluster

The effect of MEK inhibition showed in figure 3c was evaluated over CAF subtypes transcriptome by differential gene expression analysis between treated and vehicle cells with Seurat FindMarkers function^93^. The results were filtered according to pvalue<0.05 and 0.25<log2FoldChange<-0.25. To assess the purity of the CAF cluster, we extrapolated the fibroblast cells from the integrated atlas and performed subclustering with FindCluster function of Seurat ^93^ using a resolution of 0.4. Resolution value was decided after evaluation with clustree ^98^. The unique features of the fibroblast subclusters were evaluated by marker genes expression. Of the 7 subclusters, 4 were annotated as CAFs (fibroblast_c1 = 350 cells, fibroblast_c2 = 152 cells, fibroblast_c3 = 680 cells, fibroblast_c4 = 710 cells) due to expression of both subtype specific signature genes ^6^ and panCAF genes (*Pdpn*, *Fap*, *Acta2, Pdgfra*). One subcluster was annotated as cycling (140 cells) and the two remaining subclusters were annotated as EMT-like (61 cells) and Myeloid-like (53 cells) based on the high expression level of *Ptprc, Pecam1,* and *Epcam* for EMT-like cells and *Itgam, Ptprc,* and *Adgre1* for Myeloid-like cells. The 3 non-CAFs clusters were filtered out before the subsequent analysis. Differentiation analysis of CAF subtypes was performed with CytoTRACE package ^44^ using default parameters. To perform velocity analysis, the matrices of spliced and unspliced RNA counts were obtained from raw data using velocyto pipeline ^46^ under default parameters. The resulting matrices were uploaded in Seurat objects selecting cells from the previously annotated CAFs subclusters. The objects were integrated with Seurat integration pipeline by timepoints, regressing out cell cycle effect. Computation of velocity and velocity pseudotime values was performed through velociraptor R package ^45^, using default parameters and precomputed PCA values. The sMEK signature was derived by filtering the results of differential gene expression performed with Seurat function FindMarkers ^93^ between vehicle and MEKi treated CAFs subclusters of the 2 days timepoint, filtering according to pvalue<0.05 and log2FoldChange<-0.25. Signature enrichment score were computed on fibroblast compartment using UCell R package ^95^. CAFs of the vehicle samples were classified as sMEK^high^ CAF when their signature enrichment score was above the third quartile of the score’s distribution. The same value was then used on treated samples as threshold for classification of cells in sMEK high and low subclasses. Enrichment pathway analysis on the sMEK^high^ and sMEK^low^ CAFs subclass was performed with both PROGENy ^49^ and fgsea package. As input was used the ordered gene list from differential gene expression analysis performed with FindMarkers ^93^ function and pathways from HALLMARK, REACTOME, and GO collections from msigDB ^96,97^.

### Additional mouse scRNA-seq data

Single cell RNA-seq normalized counts of fibroblasts enriched dataset from Elyada et al. ^6^ were downloaded from GEO (GSE129455). The data were imported and managed with Seurat ^93^. sMEK signature enrichment score was computed on dataset cells with UCell package ^95^. Subtyping of the fibroblasts cluster was performed as described in the subtyping section of our mouse single cell RNA sequencing.

### Human single cell RNA sequencing datasets

Single cell RNA-seq dataset from Peng et al. ^51^ (primary PDAC = 24, ncells = 41964) was downloaded from Genome Sequence Archive (GSA) Project PRJCA001063 with dataset annotation metadata. The data were pre-processed using Seurat ^99^ for quality control and filtering (percent_mt_max = 20, nFeature_min = 500, nCount_min = 500, nCount_max = 50000). Pan-cancer dataset from Luo et al.^56^ (primary tumour = 148, ncells = 494610) was downloaded from Gene expression Omnibus (accession No. GSE210347) with annotation metadata^6,19^.^6,19^ sMEK signature enrichment score was computed on dataset cells with UCell package ^95^. Subtyping of the epithelial and fibroblast clusters was performed as described in the subtyping section of mouse single cell RNA sequencing.

## ACKNOWLEDGMENTS

We gratefully acknowledge the Centro Piattaforme Tecnologiche (CPT - University of Verona, Verona, Italy) for granting access to the genomic facility of University of Verona. We would like to thank Dr. T. Jacks, Dr. D. Tuveson, Dr. A. Berns and Dr. R.M. Schmid for providing transgenic animals. We are thankful to the West German Biobank Essen for infrastructural support.

## AUTHOR CONTRIBUTIONS

LV, DP, PFC, JS, and VC designed the research; LV, PFC, RF, CV, DF, EF, FL, SDA, CC, DB and MB performed experiments; DP, PD and CN analysed omics data and generated displays; RTL, CL, and AS collected and characterized human tissue samples; GB supervised *in vitro* experiments with mPSCs; VC and JTS acquired funding; LV, DP, JS, and VC wrote the manuscript; PFC, JS, and VC supervised the study. All authors approved the final version of the manuscript.

## DATA AVAILABILITY

All data relevant to this study are available upon request. All processed data generated for this study are provided in the Supplementary tables. RNA-seq, and scRNA-seq data will be made available at time of publication (or on request of the reviewers).

