## Supplementary figures for "Differential Activity of MAPK signalling Defines Fibroblast Subtypes in Pancreatic Cancer"

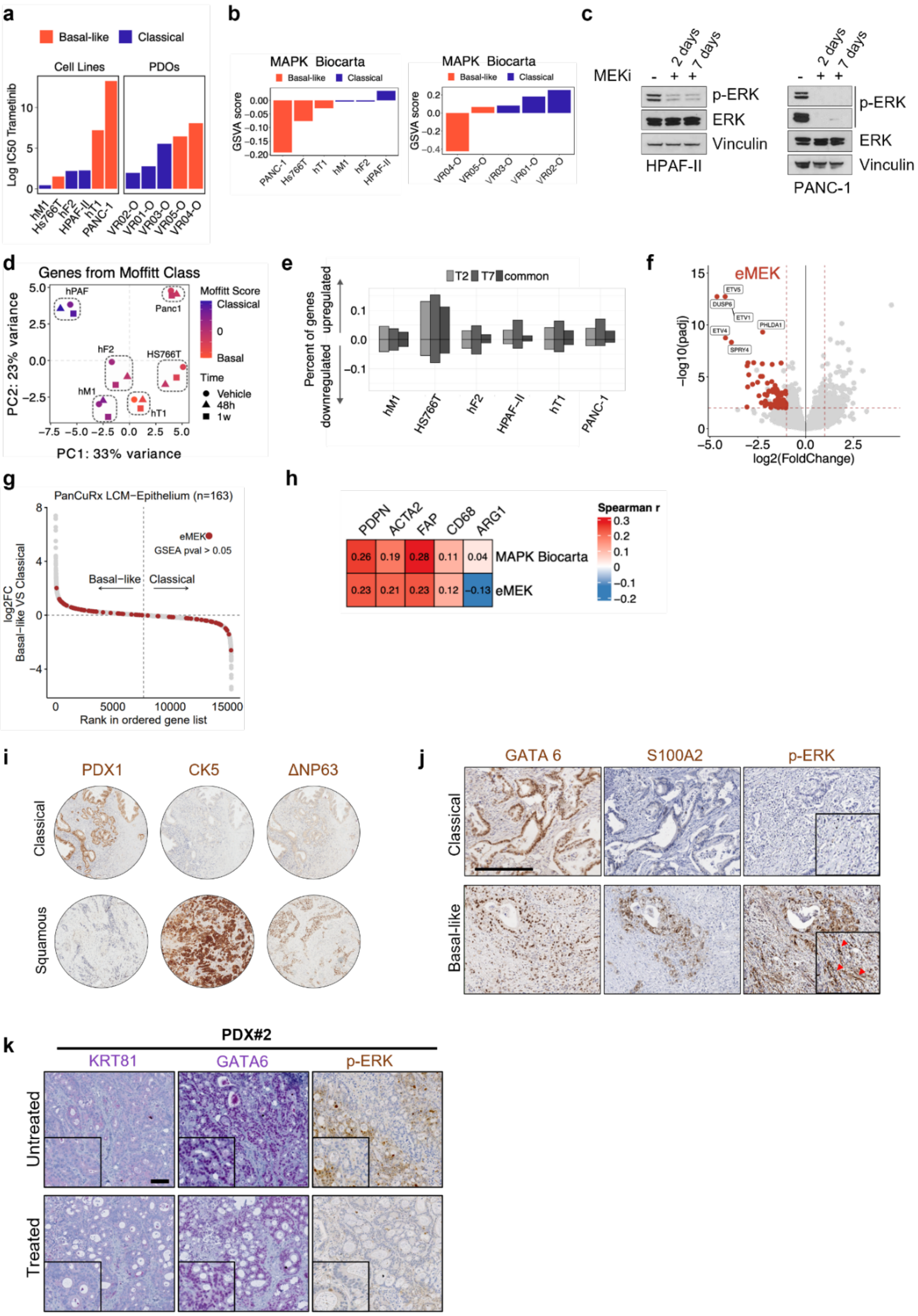


**Supplementary Figure 1: Short-term MEK inhibition does not affect PDAC epithelial subtype.** **a** Barplot showing the Log IC50 for MEKi of 6 human cell lines and 5 human organoids classified as either classical or basal-like according to Moffitt's subtypes[19]. **b** Waterfall plot showing the GSVA score for the MAPK Biocarta signature of 6 human cell lines and 5 human organoids classified according to Moffitt's classification [19] and sorted by value, in ascending order. **c** Immunoblot of p-ERK and total ERK in whole-cell lysate from HPAF-II and PANC-1 treated with either vehicle or MEKi for 2 and 7 days. Vinculin was used as loading control. **d** Principal component analysis of RNA-seq data of 6 human cell lines treated with either vehicle or MEKi for 2 or 7 days, performed on Moffitt's signatures genes. The colour scale represents the score of Moffitt's subtypes[19]. **e** Barplot showing the percentage of genes up- and down-regulated in 6 cell lines upon treatment with MEKi for 2 or 7 days. **f** Volcano plot representing the genes down- and up-regulated upon treatment with MEKi for 2 days *in vitro.* The red dots are the genes of the eMEK signature (n=89). Highlighted some of the genes with log2FC expression < -2 and adjusted $p < 0.01$. **g** Scatter plot showing the distribution of genes from eMEK signature ordered according to Moffitt's subtypes[19] in PanCuRx [17]cohort. **h** Heatmap showing Spearman's correlation between gene signatures and selected gene markers for fibroblasts and macrophages. All boxes, $p < 0.001$. **i** Images of immunohistochemistry for PDX1, CK5, and ΔNP63 of classical ($PDX1^{+}CK5^{-}\Delta NP63^{-}$) and squamous ($PDX1^{-}CK5^{+}\Delta NP63^{+}$) tumours on human PDAC tissue microarrays. **j** Images of immunohistochemistry for GATA6, S100A2, and p-ERK on classical and basal-like human PDAC tissues. Inserts showed 2X magnification of selected areas. Scale bar, 200 μm. Red arrows indicate p-ERK in stroma. **k** Representative images of immunohistochemistry for KRT81, GATA6 and p-ERK on tissues from a patient derived xenograft (PDX#2) tissue showing no change in tumour subtype upon treatment with MEKi (n=10). Scale bar, 200 μm. Inserts showed 2X magnification of selected areas.

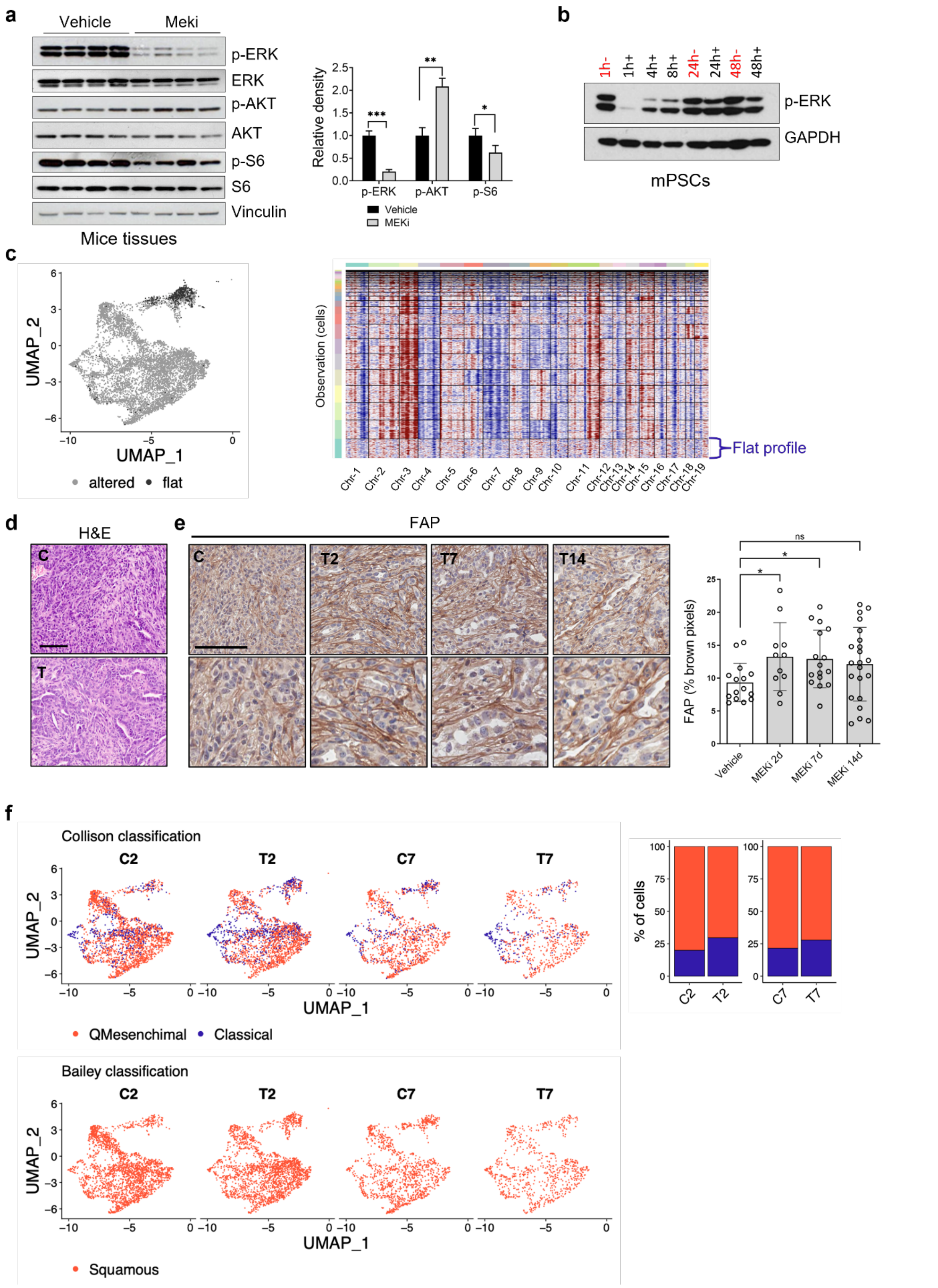


**Supplementary Figure 2: MEK inhibition is efficient at 2 days of treatment and increases the stroma content *in vivo*. a** Immunoblot of p-ERK, total ERK, p-AKT, total AKT, p-S6, and total S6 in whole-tumour lysate from mice treated for 2 days either with vehicle (n=4 mice) or MEK inhibitor 1 mg/kg (n=4 mice). Vinculin was used as loading control. Quantification of changes in the phosphorylated levels of proteins (p-ERK, p-AKT, and p-S6) is provided in the barplot on the right. ***, $p < 0.001$; **, $p < 0.01$; and *, $p < 0.05$ by Student t test (two-sided). **b** Immunoblot of p-ERK in whole-cell lysate from mPSCs treated with either vehicle (red) or MEKi at different timepoints. Vinculin was used as loading control. **c** Copy number analysis of the epithelial cluster. On the left, UMAP plot showing epithelial cells coloured according to copy number profile (grey, altered; black, flat). On the right, a representative heatmap showing inferred amplifications (red) and deletions (blue) for the sample T7 (mice treated for 7 days with MEKi) from transcriptomic data for each cell (rows) over the genomic positions (columns). Horizontal black line show cells hierarchical clustering based on CNV profile. **d** Representative haematoxylin and eosin staining of tumour tissues from vehicle (C) and MEKi (T) treated mice. Scale bar, 100 μm. **e** On the left, representative images of immunohistochemistry for FAP of tumour tissues from vehicle and MEKi treated mice at different timepoints. Scale bar, 100 μm. Lower panels show 2X magnifications of select areas. On the right, barplot showing FAP quantification provided as percentage of brown pixel per FOV. A minimum of 10 fields for each condition have been analysed. *$p < 0.05$, ns as not significant as determined by Student's t test (two-sided). Error bars show standard deviation. **f** UMAP plots showing cells of the epithelial cluster of vehicle (C) and MEKi (T) treated mice for 2 and 7 days, classified according to Collisson [18] and Bailey [15] molecular subtypes. On the right, barplot showing the percentage of cells of the epithelial cluster of mice treated either with vehicle and MEKi for 2 and 7 days, classified as classical and quasi-mesenchymal according to Collisson classification [18].

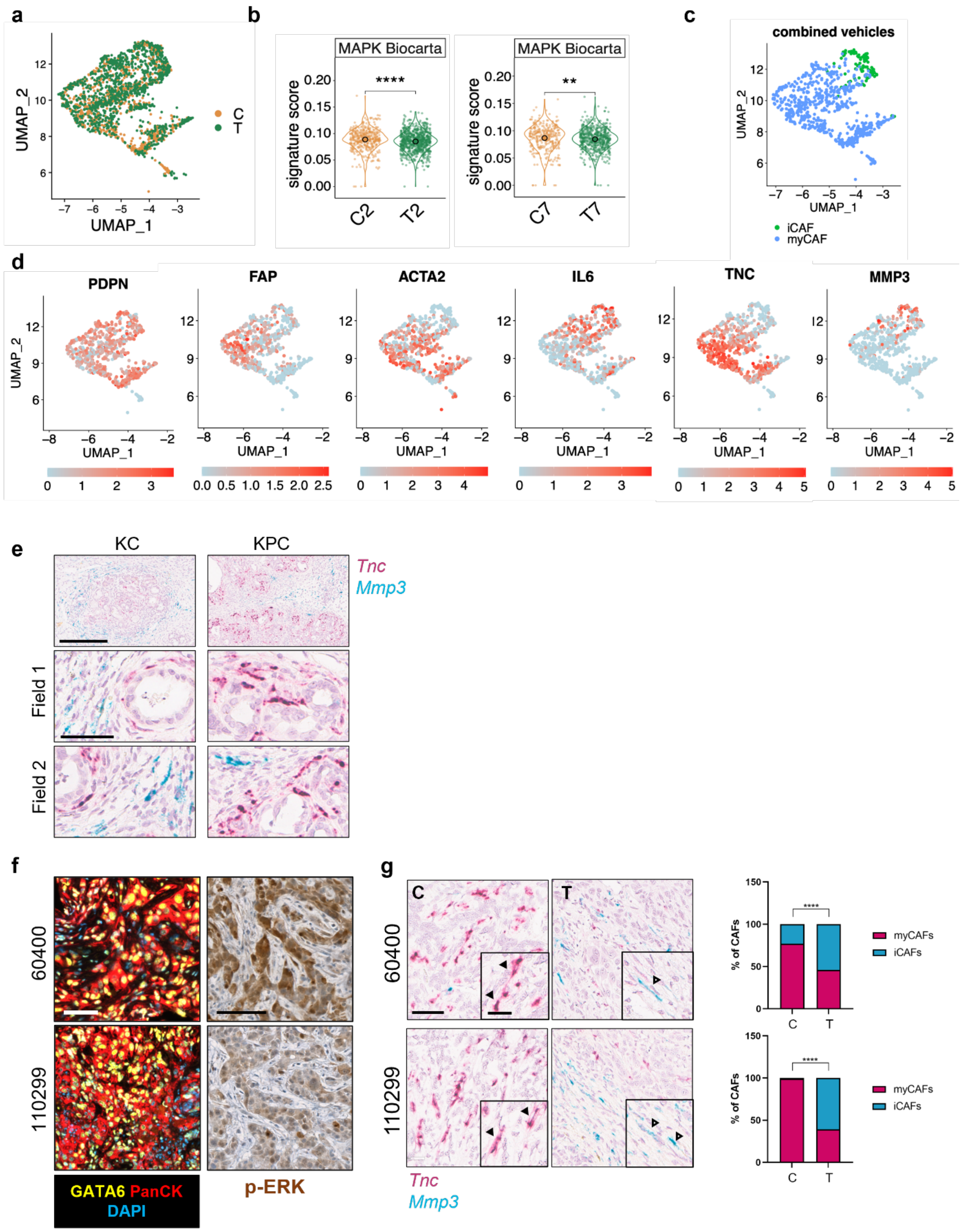


**Supplementary Figure 3: Identification of reliable markers for iCAFs and myCAFs from scRNA-seq data of mouse PDAC tumours. a** UMAP plot of cells from the fibroblast cluster of integrated samples from mice treated with either vehicle or MEKi. **b** Violin plots representing the score value for MAPK Biocarta signature of cells of the fibroblast cluster from mice treated with either vehicle or MEKi for 2 and 7 days. ****$p < 0.0001$; **$p < 0.01$ as determined by Wilcoxon test. **c** UMAP plot of cells from the fibroblast cluster from vehicle treated mice displaying the CAFs subtype for each cell according to Elyada's subtypes[6]. **d** UMAP plot of cells from the fibroblast cluster displaying the normalized expression for selected markers (*Pdpn*, *Fap*, *Acta2, Il6, Tnc,* and *Mmp3*) for each cell from vehicle treated mice. **e** Representative images of *in situ* hybridization showing expression of *Tnc* (myCAFs) and *Mmp3* (iCAFs) genes in tissues from KC and KPC mice. Scale bar, 300 μm. Lower panels show magnification of 2 selected fields. Scale bar, 60 μm. **f** Representative images of multiplex immunofluorescence performed on murine PDAC tissues derived from orthotopic transplantation of classical murine cell lines. Scale bar, 50 μm. On the right, representative images of immunohistochemistry for p-ERK performed on murine PDAC tissues derived from orthotopic transplantation of classical murine cell lines. Scale bar, 100 μm. **g** Representative images of *in situ* hybridization showing expression of *Tnc* and *Mmp3* genes on classical PDAC tissues from mice treated with either vehicle or MEKi. Scale bar, 60 μm. Insert showed a 2X magnification of selected areas. Black arrowheads indicate myCAFs, while white arrowheads point iCAFs. The quantification is provided on the right as barplot displaying the percentage of iCAFs and myCAFs content in the entire tissue slide. ****$p < 0.0001$ as determined by Student's t test (two-sided).

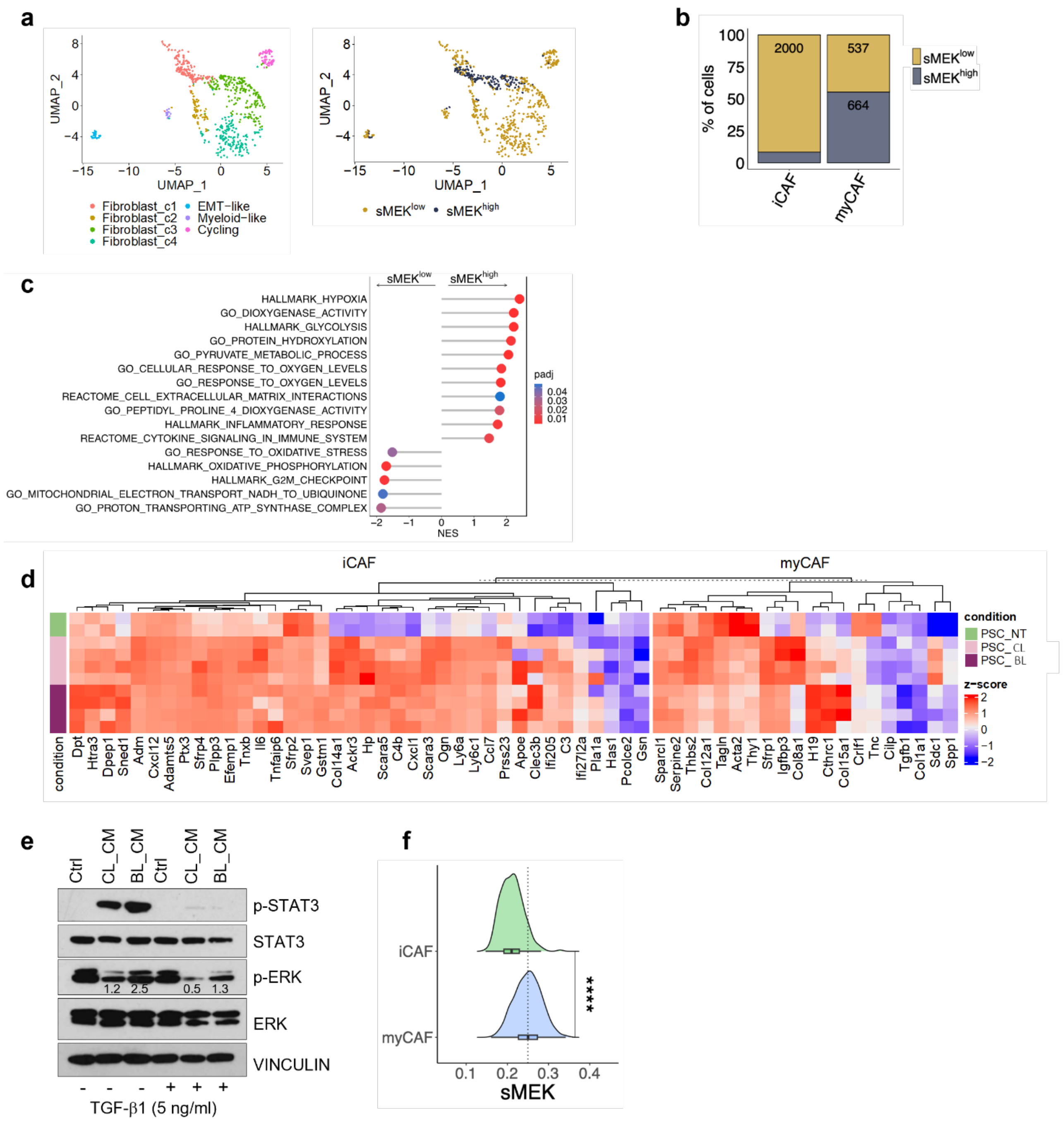


**Supplementary Figure 4: Identification of a gene expression signature of CAFs with elevated MAPK activity. a** On the left, UMAP plot of fibroblast compartment after subclustering. Different cell type clusters are colour-coded. On the right, UMAP plot of cells from the fibroblast cluster stratified according to sMEK signature from mice treated with vehicle for 2 and 7 days. **b** Barplot displaying the percentage of cells of the fibroblast cluster from the dataset of Elyada et al.[6] showing elevated MAPK activity ($sMEK^{high}$) and classified as either myCAFs or iCAFs. **c** Enrichment analysis of significantly up-regulated pathways in $sMEK^{high}$ and $sMEK^{low}$ fibroblasts. **d** Heatmap showing changes in the expression of the genes in the myCAFs and iCAFs signatures from Elyada et al.[6] in mPSCs treated with conditioned media from classical and basal-like cell lines for 24 hours. Z-scores derived from DESeq2 vst transformed counts. **e** Immunoblot analyses of p-ERK, total ERK, p-STAT3 and total STAT3 in whole-cell lysate from mPSCs either untreated or pre-treated with TGF-β1 (5 ng/ml) for 48 hours and additionally treated with conditioned media from a classical and a basal-like cell line for 1 hour. Vinculin was used as loading control. The fold change relative to control is reported as numeric values on the blot. **f** Density plot for the sMEK signature score values over the activated fibroblasts from dataset from Peng et al.[51]. Activated fibroblasts were classified as either myCAFs or iCAFs according to Elyada' et al. [6]. ****p $<$ 0.0001 as determined by Student's t test (two-sided).

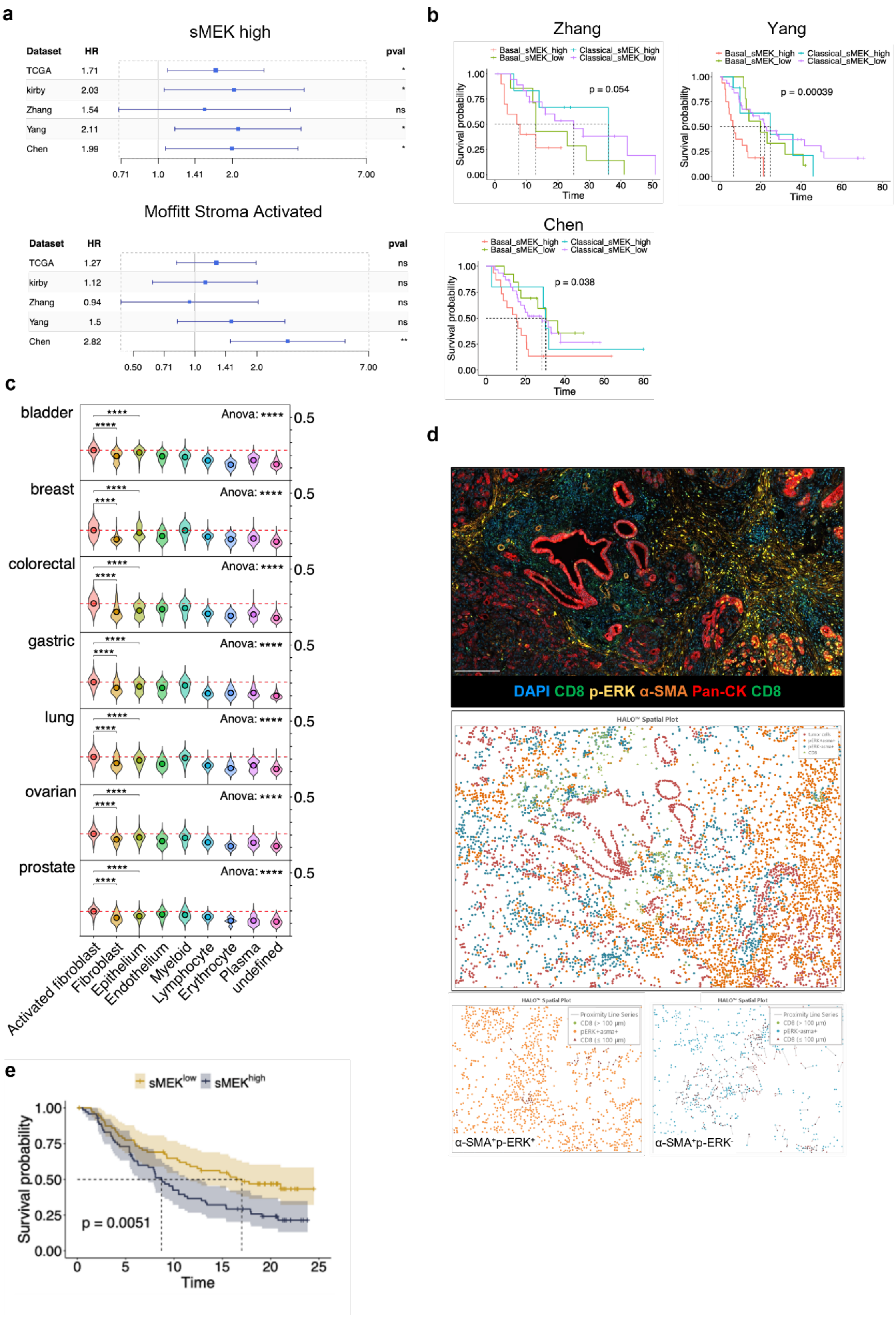


**Supplementary Figure 5: The sMEK signature can be found across several human cancer types.** **a** Forest plots showing the hazard ratio for sMEK (upper panel) and stroma activated signature from Moffitt et al.[19] (lower panel) in 5 indicated of human PDAC cohorts[16,52-55].*p<0.05; **p < 0.01; ns as not significant as determined by Wilcoxon test. **b** Combined Kaplan-Meier survival analysis comparing the survival probability of patients from 3 PDAC cohort [52,54,55] stratified according to the expression of sMEK signature and Moffitt neoplastic cells subtypes[19]. p, Log-rank (Mantel–Cox) test. **c** Violin plots representing the score values for sMEK signature of cells from different cellular type clusters in 7 cancer types from the Luo et al. dataset[56]. ****$p < 0.0001$ as determined by Anova or Wilcoxon test. **d** Spatial plots generated by Halo® Image Analysis Platform showing CD8+ cells in close proximity (100 um radial distance) of p-ERK$^{+}$α-SMA$^{+}$ and p-ERK$^{-}$ α-SMA$^{+}$ cells. **e** Kaplan–Meier plot comparing the survival probability of BLCA patients treated with immunotherapy from the IMvigor210 cohort[69]according to the expression of sMEK signature. p, Log-rank (Mantel–Cox) test.
